# Homogenization of capillary flow and oxygenation in deeper cortical layers correlates with increased oxygen extraction

**DOI:** 10.1101/450932

**Authors:** Baoqiang Li, Tatiana V. Esipova, Ikbal Sencan, Kivilcim Kilic, Buyin Fu, Michele Desjardins, Mohammad Moeini, Sreekanth Kura, Mohammad A. Yaseen, Frederic Lesage, Leif Østergaard, Anna Devor, David A. Boas, Sergei A. Vinogradov, Sava Sakadžić

## Abstract

Our understanding of how capillary blood flow and oxygen distribute across cortical layers to meet the local metabolic demand is incomplete. We addressed this question by using two-photon imaging of microvascular oxygen partial pressure (PO_2_) and flow in the whisker barrel cortex in awake mice at rest. Our measurements in layers I-V show that the capillary red-blood-cell flux and oxygenation heterogeneity, and the intracapillary resistance to oxygen delivery, all decrease with depth, reaching a minimum around layer IV, while the depth-dependent oxygen extraction fraction is increased in layer IV, where oxygen demand is presumably the highest. Our findings suggest that homogenization of physiological observables relevant to oxygen transport to tissue is an important part of the microvascular network adaptation to a local brain metabolism increase. These results will inform the biophysical models of layer-specific cerebral oxygen delivery and consumption and improve our understanding of diseases that affect the cerebral microcirculation.

**IMPACT STATEMENT:** Homogenization of cortical capillary blood flow and oxygenation underpins an important mechanism, by which the microvascular network adapts to an increase in the local brain oxidative metabolism.

## INTRODUCTION

Normal brain functioning is critically dependent on the adequate and uninterrupted supply of oxygen to brain tissue (Attwell and Laughlin, 2001; Raichle and Gusnard, 2002; Raichle et al., 2001). Significant efforts have been made over the years to investigate the regulation of cerebral blood flow (CBF), including auto-regulation of CBF in response to the changes in cerebral perfusion pressure (Aaslid et al., 1989; Ob et al., 1990) and the processes underlying neurovascular coupling (Anenberg et al., 2015; Cai et al., 2018; Girouard and Iadecola, 2006; Iordanova et al., 2015; Vazquez et al., 2010). Furthermore, neuronal and microvascular densities vary greatly between cortical layers (Blinder et al., 2013), suggesting a laminar variation of tissue metabolism (De Kock C. P. J. et al., 2007; Hyder et al., 2013). However, it is still not well understood *how blood flow and oxygenation are distributed through the microvascular network to support an adequate tissue oxygenation across the closely spaced, but morphologically and metabolically heterogeneous cortical layers*. Answering this question is important to improve our understanding of the normal brain physiology as well as brain diseases that affect cerebral microcirculation (Berthiaume et al., 2018; Iadecola, 2016, 2017; Pantoni, 2010; Zlokovic, 2011).

The recent development of tools for *in vivo* microvascular oxygen imaging enabled investigation of brain oxygen delivery and consumption within arteriolar, venular and capillary domains over large tissue volumes (Cao et al., 2017; Chong et al., 2015a; Hu et al., 2009; Lecoq et al., 2011; Parpaleix et al., 2013; Sakadžić et al., 2010, 2015; Vanderkooi et al., 1987; Wang et al., 2011; Yaseen et al., 2009). Distributions of microvascular blood flow and oxygen in mice at rest have been assessed in several studies using optical coherence tomography, two-photon microscopy and photoacoustic imaging (Cao et al., 2017; Chong et al., 2015b; Gutiérrez-Jiménez et al., 2018; Lecoq et al., 2011; Lyons et al., 2016; Moeini et al., 2018; Parpaleix et al., 2013; Sakadžić et al., 2011, 2014; Santisakultarm et al., 2012, 2014; Srinivasan et al., 2015). However, in most previous studies these variables have not been considered as a function of cortical depth and/or capillary branching order. Instead, measurements of intravascular oxygen concentrations were performed at limited cortical depths and typically under anesthesia, which could have significantly affected both brain CBF and metabolism (Alkire et al., 1999; Goldberg et al., 1966), or using samples of limited size that might prevent detailed depth-dependent mapping of microvascular blood flow and oxygen with sufficient statistical power (Lecoq et al., 2011; Lyons et al., 2016; Parpaleix et al., 2013).

To this end, in the present work we applied two-photon phosphorescence lifetime microscopy to measure the absolute intravascular partial pressure of oxygen (PO_2_) in a large number of arterioles, venules and capillaries, as well as the erythrocyte-associated transients of capillary PO_2_ (EATs), capillary red blood cell (RBC) flux and microvascular structure as a function of cortical depth within the range of 0-600 µm. We used a new phosphorescent oxygen probe, based on a recently developed Pt tetraarylphthalimidoporphyrin (PtTAPIP) (Esipova et al., 2017). Compared to its predecessor PtP-C343 (Finikova et al., 2008), the new probe exhibits longer excitation and emission wavelength maxima, higher quantum yield and a larger two-photon absorption cross-section, facilitating simultaneous mapping of microvascular PO_2_ and capillary RBC flux in cortical layers I-V in mice and making it possible to significantly extend both the sample size and imaging depth in comparison to the previous studies (Lecoq et al., 2011; Lyons et al., 2016; Parpaleix et al., 2013; Sakadžić et al., 2010, 2014). Moreover, our measurements were performed in head-restrained awake mice, and thus were free of the confounding effects of anesthesia on neuronal activity, CBF and brain metabolism. We found that in the whisker barrel cortex in awake mice the oxygen extraction fraction (OEF), measured at different cortical depths (i.e. depth-dependent OEF), reached its maximum at the depths of 320-450 µm, corresponding to cortical layer IV, where the neuronal and capillary densities are the largest, and, presumably, oxygen consumption is the highest. The increased OEF was accompanied by homogenization of the capillary PO_2_ and RBC flux, as well as by a decrease in the intracapillary resistance to oxygen delivery (inferred from the EATs amplitude). These experimental results enabled quantification of parameters of importance for oxygen transport to tissue and put forward a potential mechanism, by which the microvascular networks at rest may adapt to the heterogeneous metabolic demands in different cortical layers. We anticipate that our results will improve our understanding of the normal brain function as well as of diseases that affect the cerebral microcirculation (Girouard and Iadecola, 2006; Iadecola, 2016; Müller et al., 2017; Pantoni, 2010; Wardlaw et al., 2013; Zlokovic, 2011). In addition, detailed knowledge of the distributions of capillary RBC flux and oxygenation at different depths should inform the next generation of biophysical models of the layer-specific oxygen delivery and consumption (Gagnon et al., 2016; Mintun et al., 2001; Secomb et al., 2000).

## RESULTS

### Accumulated oxygen extraction fraction increases in deeper cortical layers

We used a home-built two-photon microscope (Sakadžić et al., 2010; Yaseen et al., 2015) (Fig. 1a) to measure the resting intravascular PO_2_ in the whisker barrel cortex in head-restrained awake mice through a chronic cranial window. PO_2_ imaging was performed within a 500×500 µm^2^ field of view (FOV) down to 600 µm below the cortical surface. At each imaging depth, we selected the scan points for measuring PO_2_ inside the microvascular segments (one point per segment), including arterioles, venules and capillaries (Fig. 1f). Point-based acquisition of PO_2_ was conducted plane-by-plane from the cortical surface down to the cortical depth of 600 µm with inter-plane separation ≤50 µm. The intravascular Mean-PO_2_ was measured in all diving arterioles, venules and in the majority of branching arterioles, venules and capillaries (6,544 vascular segments across n = 15 mice) within the FOV. In addition, capillary RBC flux, speed and hematocrit, as well as RBC-PO_2_, InterRBC-PO_2_, and EATs (please see the METHODS section for the details) were calculated in a large subset of capillary segments (978 capillaries across n = 15 mice). In each mouse, the measurements were grouped by the cortical layer and then averaged. Subsequently, the average measurements for each cortical layer were averaged over animals (please see the METHODS section for the details). Histograms of capillary Mean-PO_2_, RBC flux, hematocrit and speed are presented in Supplementary Fig. 2.

**Figure 1.**
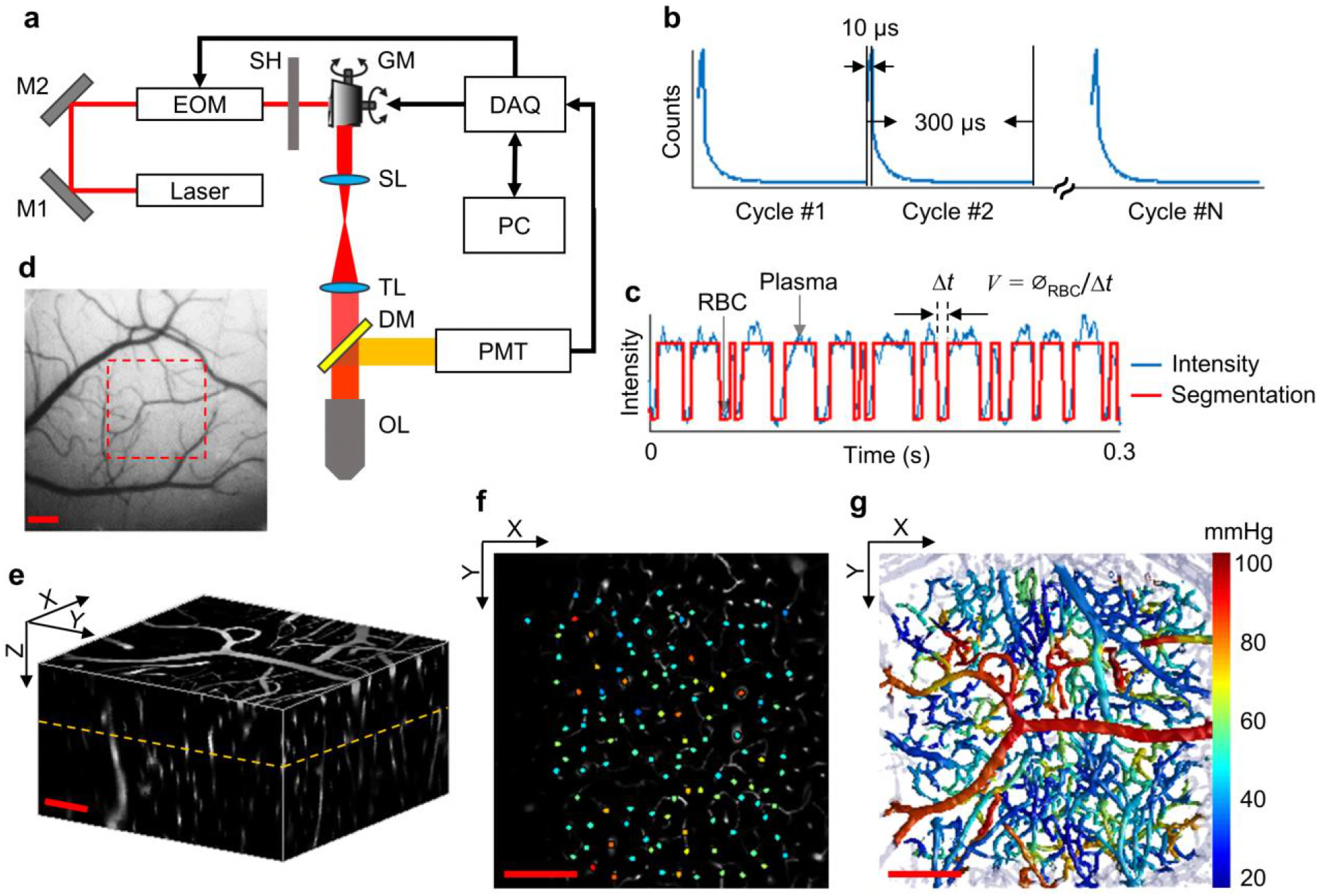
Experimental setup and data acquisition protocol. **a.** Schematic of our home-built two-photon microscope. The components are abbreviated as: mirror (M), electro-optic modulator (EOM), shutter (SH), galvo mirrors (GM), scan lens (SL), tube lens (TL), dichroic mirror (DM), objective lens (OL), and photomultiplier tube (PMT). **b.** Illustrative example of the phosphorescence decays for PO_2_ recording at a single location. Each 300-μs-long cycle includes a 10-μs-long EOM-gated excitation, followed by a 290-μs-long detection of phosphorescence decay. **c.** A representative phosphorescence intensity time course during PO_2_ recording at a single location (blue curve). Each point in the time course represents the sum of the photon counts acquired during one 300-μs-long excitation/decay cycle in **b**. The red curve represents the binary segmented time course, with valleys and peaks representing RBC and blood plasma passages through the focal volume, respectively. **d.** An image of the brain surface vasculature, taken through the chronic cranial window using a CCD camera. **e.** The 3D representation of a Sulforhodamine-B labeled cortical microvasculature imaged over the region of interest outlined by the red dashed square in **d**. **f.** PO_2_ measurements inside the microvascular segments at the imaging plane outlined by the orange dashed line in **e**. PO_2_ values (in mmHg, color-coded) were spatially co-registered with the microvascular angiogram. **g.** Composite image shows the top view of the 3D projection of the PO_2_ distribution in the microvascular network. The color bar serves for panels **f** and **g**. Scale bars: 200 µm.

The average PO_2_ values in the diving arterioles and surfacing venules across cortical layers I-V are shown in Fig. 2a. The PO_2_ in the diving arterioles decreased from 99±4 mmHg in layer I to 84±3 mmHg in layer V; while the PO_2_ in the surfacing venules exhibited a small increase starting from 43±3 mmHg in layer V to 49±4 mmHg in layer I. Similar trends were observed in the SO_2_ values (Fig. 2b), but the amplitudes of SO_2_ changes were different from those of PO_2_ due to the sigmoidal shape of the oxygen-hemoglobin dissociation curve (Uchida et al., 1998). The SO_2_ in the diving arterioles decreased slightly from 90.9±0.8 % in layer I to 87.1±1.0 % in layer V (ΔSO_2,A_ = 3.8±1.2 %), while the SO_2_ change in the surfacing venules was larger, from 53.5±1.0 % in layer V to 61.8±0.8 % in layer I (ΔSO_2,V_ = 8.3±1.1 %). Consequently, the layer-specific difference between the SO_2_ in the diving arterioles and that in the surfacing venules increased towards deeper layers, and the depth-dependent OEFs (DOEF) in layers IV (39.9±3.2 %) and V (38.7±3.3 %) were larger than in more superficial layers, e.g. layers I (32.0±2.7 %) and II/III (33.3±2.7 %; Fig. 2b).

**Figure 2.**
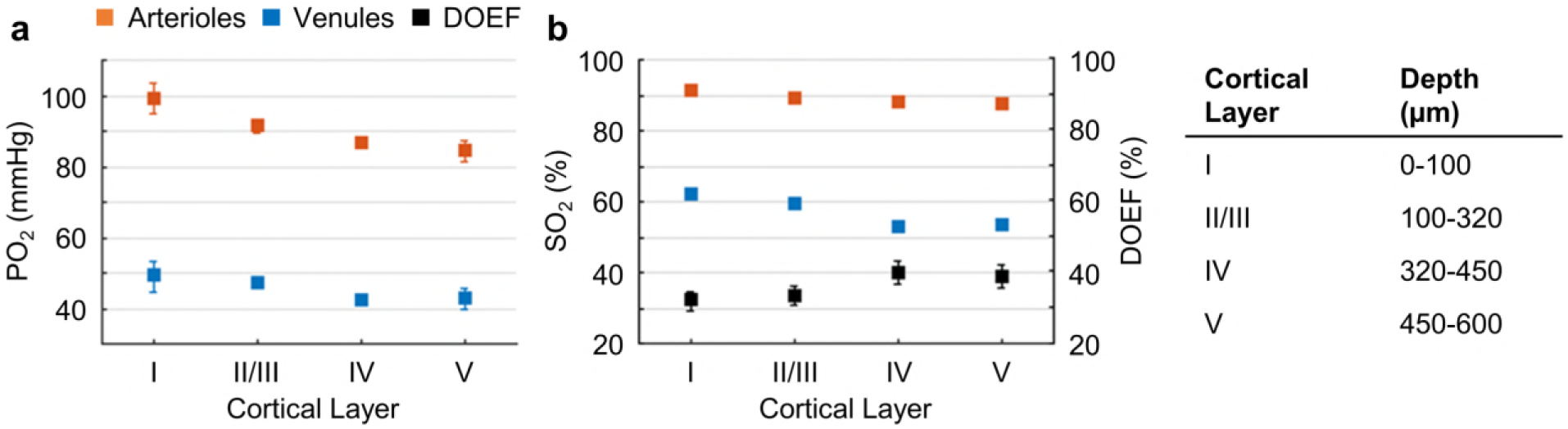
Cortical layer-dependent distributions of the arteriolar and venular intravascular PO_2_ and SO_2_. **a.** Intravascular PO_2_ in the diving arterioles (red symbols) and venules (blue symbols) across cortical layers I-V (11 arterioles, 14 venules, from n = 7 mice). **b.** SO_2_ in the diving arterioles (red symbols) and venules (blue symbols) across cortical layers I-V, and the depth-dependent OEF (DOEF, black symbols). For each diving arteriole or surfacing venule in **a** and **b**, PO_2_ was tracked from the cortical surface down to the cortical depth of 600 μm. Data are expressed as mean±SEM.

The observed increase in the DOEF with cortical depth suggests that surfacing venules received more oxygenated blood in the upper cortical layers than in the deeper layers, and that SO_2_ in the pre-venular capillaries (or “downstream capillaries”) was higher for the capillaries joining venules in the upper cortical layers than for those in the deeper layers. We revisit this observation in the later section.

### Capillary RBC flux and oxygenation homogenize in deeper cortical layers

To better understand how the distributions of the capillary RBC flow and oxygenation change in order to support the heterogeneous demand for oxygen across cortical layers, we assessed both the spatial distributions and the temporal fluctuations of the resting capillary RBC flux, speed and Mean-PO_2_ in layers I-V. The spatial distributions for both the mean value and heterogeneity (quantified by STD and CV across capillaries) of capillary RBC flux, speed and Mean-PO_2_ in cortical layers I-V are shown in Fig. 3. The RBC flux in layers IV (36±4 RBC/s) and V (38±6 RBC/s) were slightly lower than in layers I (41±2 RBC/s) and II/III (41±1 RBC/s; Fig. 3a). The decrease in the RBC flux in the deeper layers might be due to the redistribution of RBCs over a denser capillary network, especially in layer IV (Blinder et al., 2013; Sakadžić et al., 2014), causing the RBC flux to be lower in individual capillaries. Importantly, both the STD and CV of RBC flux were lower in layers IV-V than in layers I-III, reaching a minimum in layer IV (Fig. 3b, c); and the STD and CV of RBC flux in layer IV were significantly lower than their counterparts in layer I. This result suggests that a significant homogenization of RBC flux occurred in the deeper cortical layers, which is in line with the biophysical models that predict that capillary blood flow homogenization improves the efficiency of oxygen delivery to brain tissue (Jespersen and Østergaard, 2012). Similar trends in homogenization could be observed in the distributions of RBC speed (Fig. 3d-f) and capillary Mean-PO_2_ (Fig. 3g-i), as well as in the distributions of capillary RBC-PO_2_ and SO_2_ (Supplementary Fig. 3).

**Figure 3.**
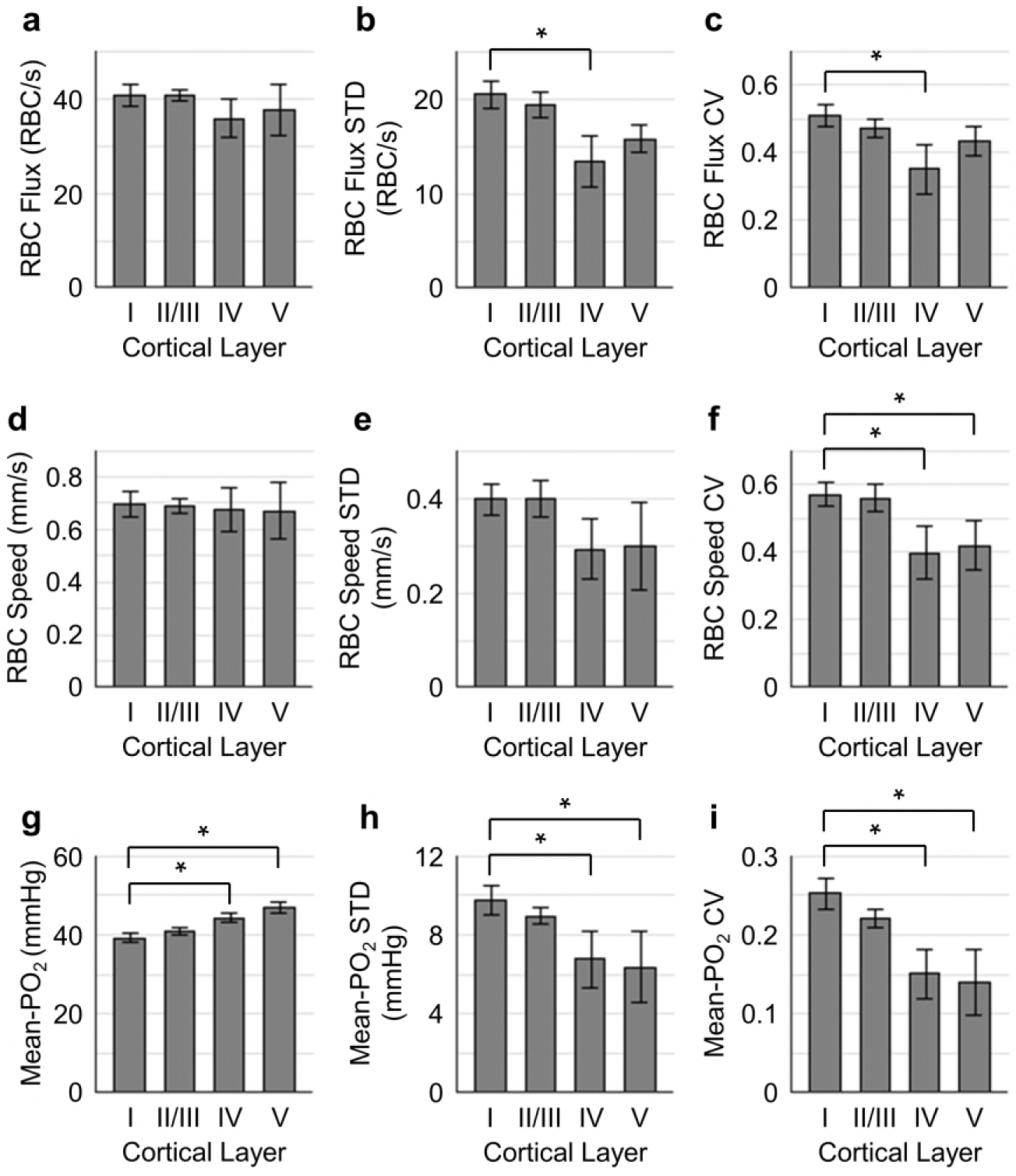
Spatial variations of capillary RBC flux, speed, and Mean-PO_2_ as a function of cortical depth. The panels **a-c**, **d-f**, and **g-i** show the dependence of the absolute values, standard deviations (STDs), and coefficients of variance (CVs) of capillary RBC flux, speed, and Mean-PO_2_ on cortical layer, respectively. The absolute values, STDs, and CVs were calculated across capillaries. The analysis in **a-i** was made with 400, 356, 118, and 104 capillaries measured in cortical layers I, II/III, IV and V, respectively, across n = 15 mice. Data are expressed as mean±SEM. The asterisk symbol (*) indicates significant difference (Student’s t-test, P<0.05).

Next, we assessed the temporal fluctuations of RBC flux, speed and Mean-PO_2_ within individual capillaries, extracted from the 9-s-long acquisitions (Fig. 4). The level of the temporal fluctuations (quantified by STD and CV) of capillary RBC flux (Fig. 4a, b) and speed (Fig. 4c, d) decreased only in layer V, while the temporal fluctuation of the capillary Mean-PO_2_ was significantly attenuated in layers IV and V (Fig. 4e, f). However, the mean amplitude of the temporal fluctuation of each observable was significantly smaller than that calculated across different capillaries (Fig. 3).

**Figure 4.**
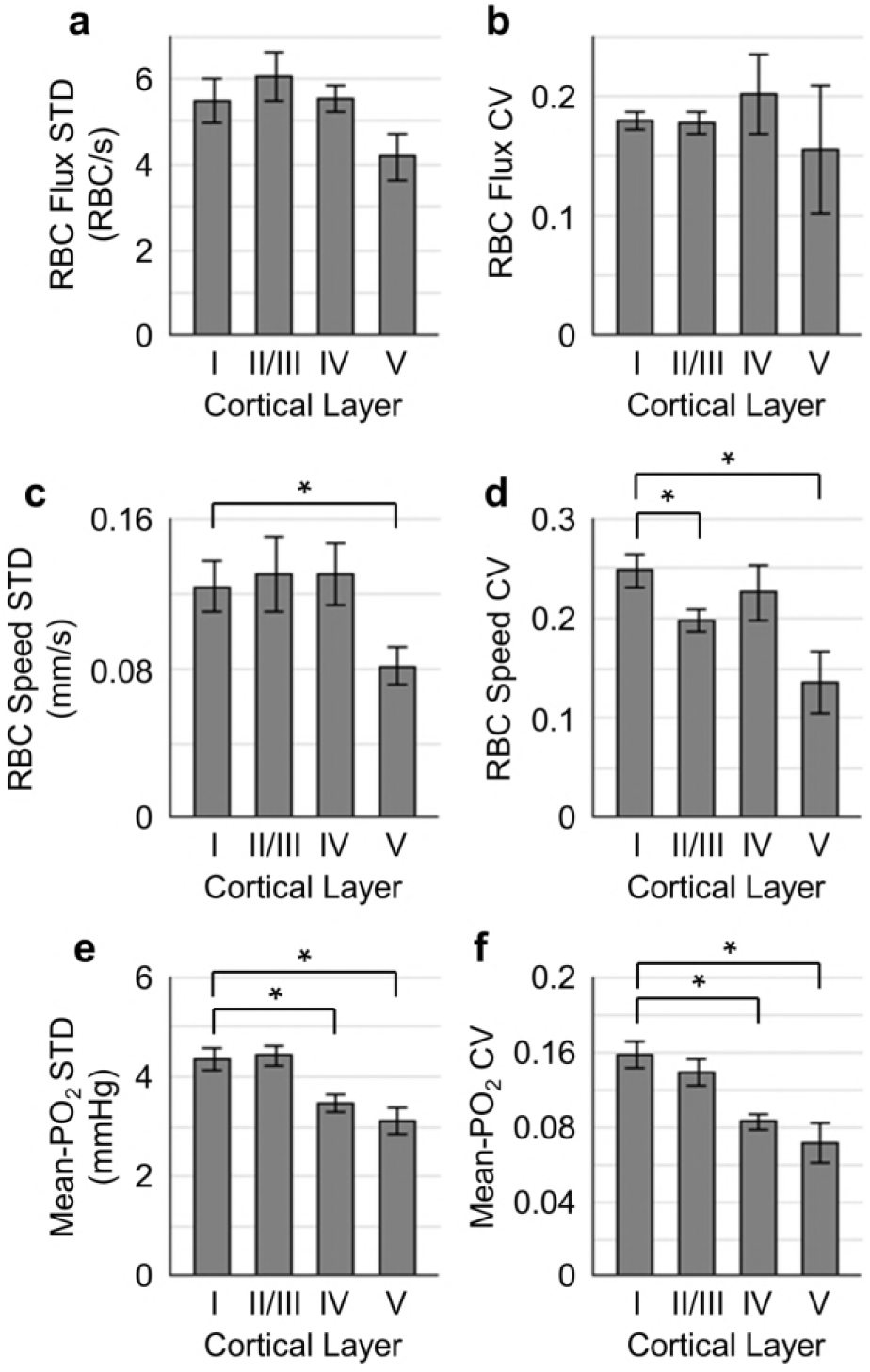
Temporal fluctuations of RBC flux, speed, and Mean-PO_2_ within individual capillaries in cortical layers I-V. Panels **a-b**, **c-d**, and **e-f** show the layer-dependent standard deviations (STDs) and coefficients of variance (CVs) of the temporal fluctuations of RBC flux, speed, and Mean-PO_2_, respectively. The STD and CV of each observable for each capillary were calculated based on the 9-s-long time course. The analysis in **a-f** was made with 130, 140, 63, and 40 samples, collected in cortical layers I, II/III, IV, and V, respectively, across n = 7 mice. Each sample corresponds to a 9-s-long, 15-time-point measurement acquired in each capillary. Data are expressed as mean±SEM. The asterisk symbol (*) indicates significant difference (Student’s t-test, P<0.05).

Importantly, our measurements revealed that the Mean-PO_2_, measured within individual capillaries as a function of time, was best correlated with RBC flux (Fig. 5). We calculated the Pearson correlations between the temporal fluctuations of RBC flux, speed, hematocrit and the temporal fluctuation of the Mean-PO_2_. All these parameters were simultaneously measured in each assessed capillary (n = 373 capillaries). The correlation coefficient *r* between the temporal fluctuations of the RBC flux and Mean-PO_2_ (median value = 0.71) was higher than between the RBC speed and Mean-PO_2_ (median value = 0.37), and between hematocrit and Mean-PO_2_ (median value = 0.29). The *r* values were converted to Fisher z values to compare for statistical difference (Diamond et al., 2006). Mean-PO_2_ was significantly stronger correlated with RBC flux than with RBC speed and hematocrit (Fig. 5c). The results of the pairwise correlations between the temporal fluctuations of capillary RBC flux, speed and hematocrit are presented in Supplementary Fig. 4.

**Figure 5.**
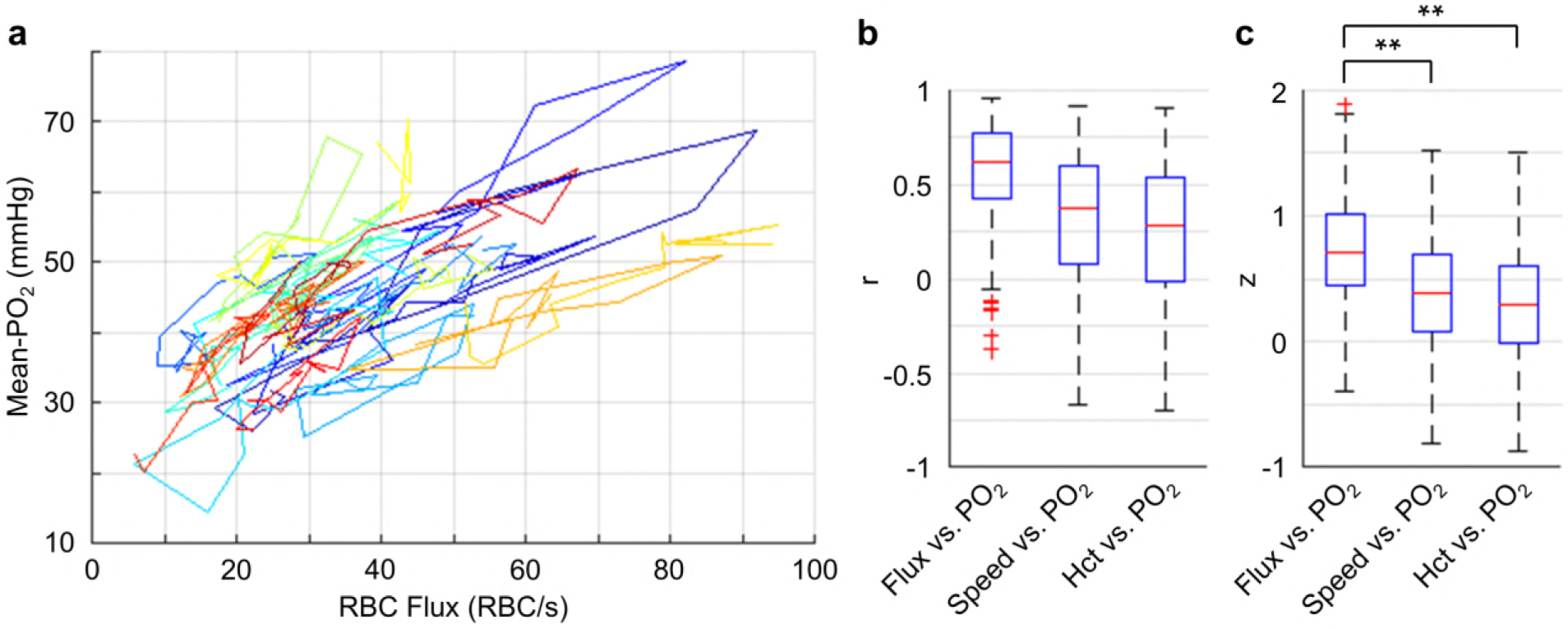
Correlations between the temporal fluctuations of capillary RBC flux, speed, hematocrit, and Mean-PO_2_. **a.** Temporal evolutions (9-s-long traces with 0.6-s steps) of RBC flux and Mean-PO_2_ from 30 representative capillaries. For each capillary, its Mean-PO_2_ as a function of RBC flux is represented by a different line color; the consecutive time points are connected to illustrate the temporal trajectory of the variation. **b.** Boxplots of the pairwise correlation coefficients (r) between the temporal fluctuations of capillary RBC flux, speed, hematocrit (Hct), and Mean-PO_2_. **c.** Boxplots of the Fisher *z* values calculated based on the *r* values in **b**. The analysis in **b** and **c** was made with 373 capillaries, collected in cortical layers I-V, across n = 7 mice. The double asterisk symbol (**) indicates significant difference (Student’s t-test, P<0.001).

Our depth-resolved measurements across capillaries revealed that the distributions of the RBC flux, speed and Mean-PO_2_ became more homogeneous in layers IV and V compared to layers I-III. This increase in the spatial homogenization of the capillary flow and oxygenation (Fig. 3) was accompanied by an increase in the depth-dependent OEF (Fig. 2). We also observed attenuation of the temporal fluctuations of these parameters in layers IV and V compared to layers I-III (Fig. 4), although with much lower amplitudes than seen in the corresponding spatial variations (Fig. 3). Finally, the time-course cross-correlation analysis revealed that the Mean-PO_2_ was best correlated with the RBC flux, as opposed to RBC speed and hematocrit (Fig. 5).

### Intracapillary resistance of oxygen transport to tissue decreases in deeper cortical layers

We now turn to the analysis of EATs that reflect capillary modulation of PO_2_ due to the passages of individual RBCs that have been observed in the peripheral and brain capillaries alike (Barker et al., 2007; Golub and Pittman, 2005; Lecoq et al., 2009). Larger EATs amplitudes have been associated with higher intracapillary resistance to oxygen transport to tissue from capillaries (Barker et al., 2007; Golub and Pittman, 2005; Hellums, 1977). Since cortical layers exhibit differences in neuronal and vascular density and possibly in oxidative metabolism, we examined whether the EATs amplitudes would differ across cortical layers. Benefitting from the improved sensitivity of the new oxygen probe, we were able to measure intracapillary longitudinal PO_2_ gradients in a larger number of capillaries (Fig. 6a; 373 capillaries across n = 7 mice) and with greater imaging depth (down to the cortical depth of 600 μm) in comparison to the previous studies (Lecoq et al., 2011; Lyons et al., 2016; Parpaleix et al., 2013). Averaging over all the assessed capillaries, PO_2_ (black curve in Fig. 6a) decreased from 55.0±0.6 mmHg at the RBC center to 30.0±7.8 mmHg, 30 μm away from the RBC center, where the PO_2_ measurements were typically associated with the capillaries with low hematocrit. However, in most capillaries the half-distance between adjacent RBCs was much smaller than 30 μm. The mean half-distance between adjacent RBCs was estimated to be ∼8 μm (denoted by the gray arrow in Fig. 6a), close to the previously reported 7 μm measured in anesthetized rats (Kleinfeld et al., 1998). Therefore, the estimated mean amplitude of EATs across all the assessed capillaries was 12.5±0.4 mmHg, significantly smaller than that in the capillaries with low hematocrit (Supplementary Fig. 5). The median values of capillary Mean-PO_2_, RBC-PO_2_, InterRBC-PO_2_, and EATs were 39.4 mmHg, 49.2 mmHg, 36.3 mmHg, and 11.7 mmHg, respectively (Fig. 6b). Interestingly, we found a significant reduction in the EATs amplitude in cortical layers IV (9.9±0.8 mmHg) and V (11.0±0.8 mmHg) compared to layers I (13.4±0.6 mmHg) and II/III (13.3±0.3 mmHg; Fig. 6c), suggesting that the intracapillary resistance to oxygen transport to tissue may be lower in deeper cortical layers, thus facilitating oxygen delivery to brain tissue in accordance with biophysical modelling (Hellums, 1977).

**Figure 6.**
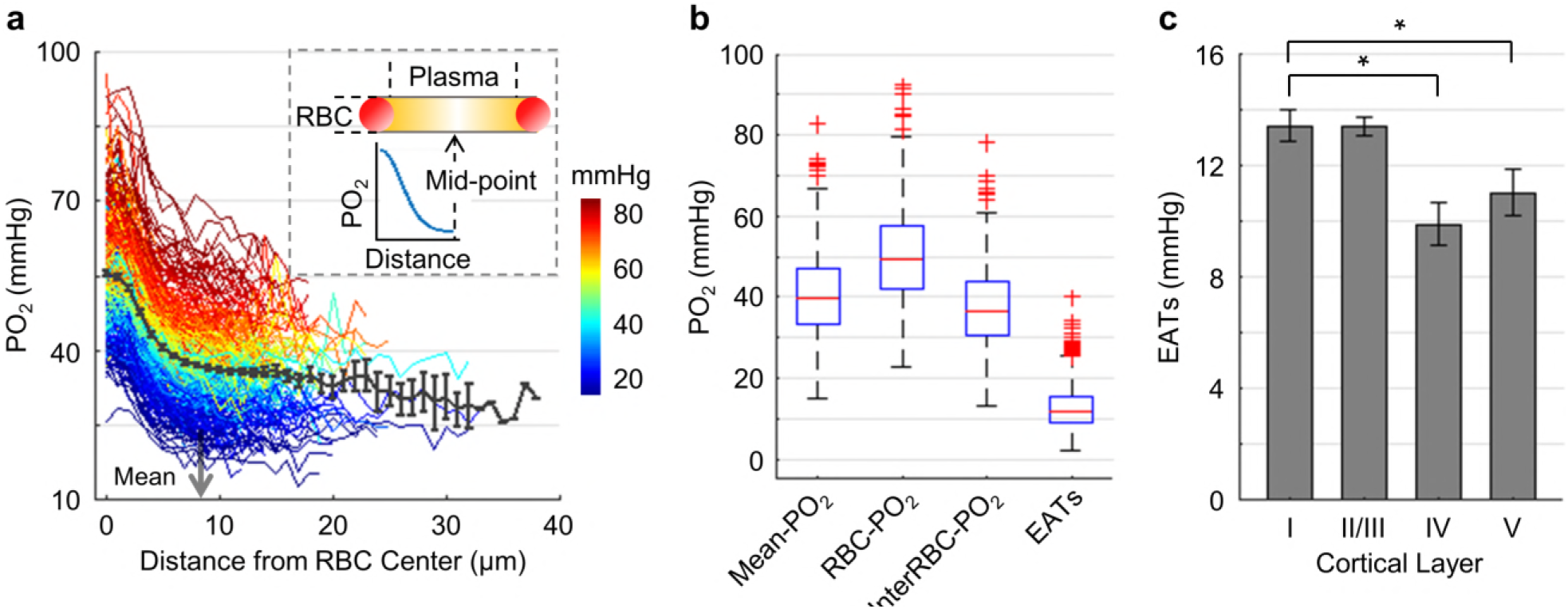
Dependence of EATs on cortical layer. **a.** Intracapillary longitudinal PO_2_ gradients. The PO_2_ gradients measured in different capillaries (373 capillaries across n = 7 mice) are color-coded based on their Mean-PO_2_ (color bar in mmHg). The black curve represents the average PO_2_ gradient. The gray arrow denotes the mean half-distance between adjacent RBCs (∼8 μm). The inset schematically illustrates the intracapillary PO_2_ gradient. **b.** Boxplots of capillary Mean-PO_2_, RBC-PO_2_, InterRBC-PO_2_, and EATs. **c.** Dependence of EATs on cortical layer. The analysis in **c** was made with 130, 140, 63, and 40 capillaries measured in cortical layers I, II/III, IV, and V, respectively, across n = 7 mice. Data are expressed as mean±SEM. The asterisk symbol (*) indicates significant difference (Student’s t-test, P<0.05).

### Low oxygen extraction along the superficial capillary paths contributes to an increase in the mean venular SO_2_ towards cortical surface

To better understand why SO_2_ in the ascending venules increased towards the cortical surface (Fig. 2), we investigated the distributions of capillary flow and oxygenation along the capillary paths in the upper cortical layers. For this analysis, capillaries in the top 300 μm of the cortex (layers I-III) were grouped based on their branching orders into two main groups: 1) ‘upstream’ capillaries with branching orders A1-A3, which are closer to the arteriolar side of the network, and 2) ‘downstream’ capillaries with branching orders V1-V3, which are closer to the venular side of the network. This grouping provides sufficient statistical power to compare the RBC flow and oxygenation parameters in the beginning and in the end of the capillary paths while avoiding labor-intensive segmentation of the microvascular angiograms.

The average values of Mean-PO_2_, SO_2_, EATs, RBC flux, speed and hematocrit from the combined A1-A3 (upstream) and V1-V3 (downstream) capillaries are presented in Fig. 7a-f. The average Mean-PO_2_ in the A1-A3 capillaries (64.4±2.4 mmHg) in layers I-III was, as expected, significantly higher than that in the V1-V3 capillaries (41.8±0.9 mmHg; Fig. 7a), suggesting that a large fraction of oxygen has been extracted along the capillary paths (i.e. from A1 to V1). Indeed, the average SO_2_ in the A1-A3 capillaries (77.6±1.7 %) was significantly higher than that in the V1-V3 capillaries (67.3±1.2 %; Fig. 7b). Provided that the average SO_2_ in the A1 capillary segments was 81.0±2.3 %, and that the SO_2_’s in the arterioles (SO_2,A_) and venules (SO_2,V_) at cortical surface were 91.0±0.8 % and 62.0±0.8 %, respectively (Fig. 2b), we estimated that the decrease in SO_2_ from the pial arterioles to the A1 capillary segments in the upper 300 μm of the cortex accounted for 34 % (or 1/3) of the total extracted oxygen. Here, the total extracted oxygen was calculated as the A-V difference in SO_2_ at the cortical surface (ΔSO_2,A-V_ = 30 %; Fig. 2b). Therefore, 66 % (or 2/3) of the oxygen extraction in awake mice took place after the arterioles. This is in contrast to our previous study in anesthetized mice, which reported that ∼50 % of the oxygen delivered to brain tissue was extracted after the arterioles (Sakadžić et al., 2014). The average V1 capillary SO_2_ in layers I-III was 65±1.9 %, higher than that in the ascending venules, both in layer I (62±0.8 %) and layer II/III (59±0.4 %; Fig. 2b). Accordingly, capillaries that feed the surfacing venules in layers I-III apparently do so with more oxygenated blood, contributing to the increase in the venular SO_2_ towards brain surface (Fig. 2b).

**Figure 7.**
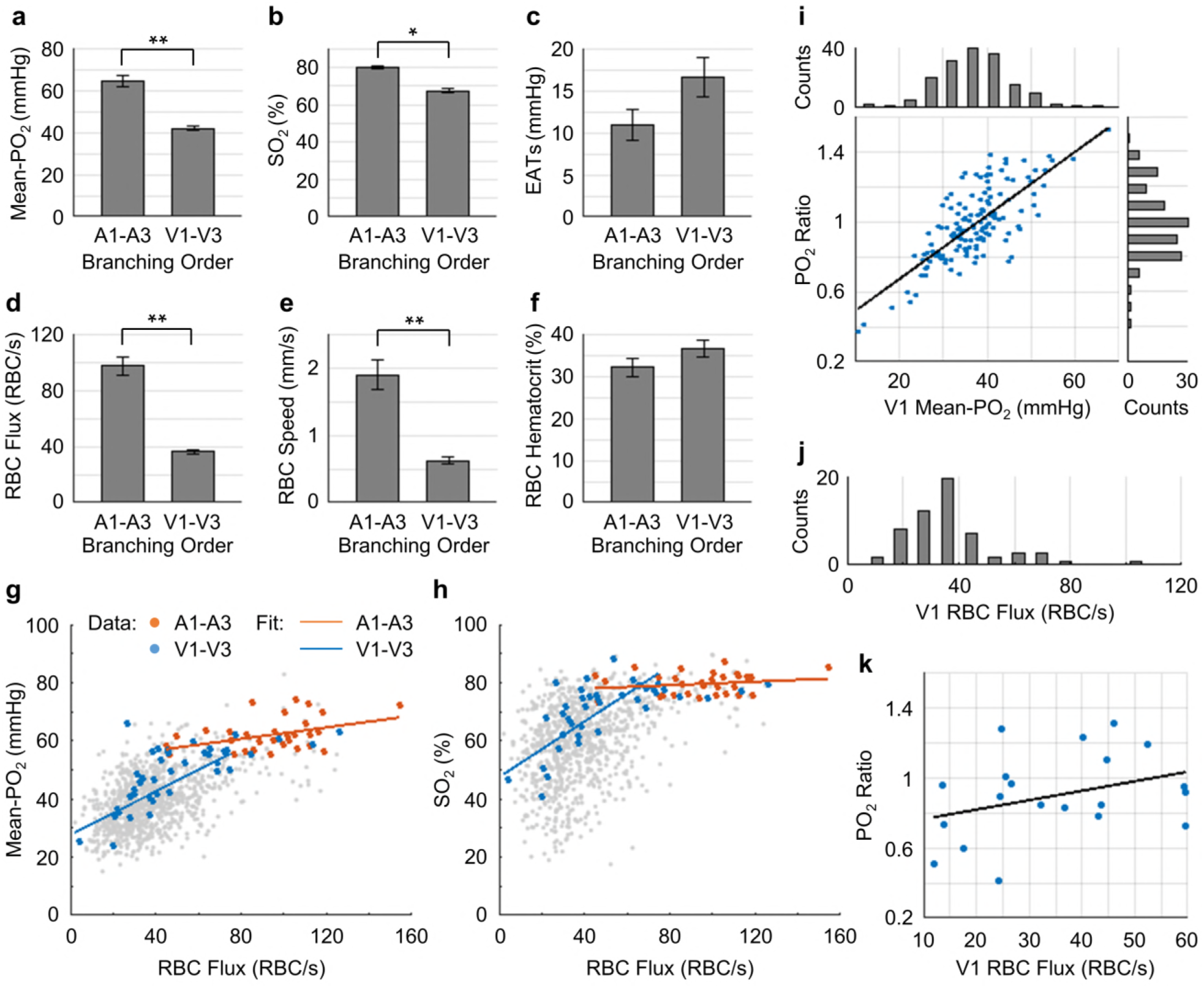
Capillary flow and oxygenation in the upstream and downstream branches. **a-f.** Average capillary Mean-PO_2_, SO_2_, EATs, RBC flux, speed, and hematocrit, in the upstream (A1-A3) and downstream (V1-V3) capillary branches, across cortical layers I-III. Data are expressed as mean±SEM. The single (*) and double (**) asterisk symbols indicate significant differences (Student’s t-test) with P<0.05 and P<0.001, respectively. **g** and **h.** Correlations between capillary RBC flux and Mean-PO_2_ (**g**) and SO_2_ (**h**). Data points and regression lines from the A1-A3, V1-V3, and all capillary segments (branching order unassigned) are color-coded red, blue, and gray, respectively. Linear regression slopes in **g**: V1-V3 slope = 0.37 mmHg·s·RBC^-1^ (R^2^ = 0.61), A1-A3 slope = 0.1 mmHg·s·RBC^-1^ (R^2^ ≈ 0.17). Linear regression slopes in **h**: V1-V3 slope = 0.50 s·RBC^-1^ (R^2^ = 0.52), A1-A3 slope = 0.03 s·RBC^-1^ (R^2^≈0.04). The analysis in **a-h** was made with 47 upstream and 50 downstream capillaries, across n = 5 mice. **i.** Correlation between the PO_2_ ratio (the V1 capillary Mean-PÜ_2_ to the adjacent PCV PO_2_) and the V1 capillary Mean-PÜ_2_. Histograms of the V1 capillary Mean-PO_2_, and PO_2_ ratio are at the top and on the right from the main panel, respectively. **j.** Histogram of the V1 capillary RBC flux. **k.** Correlation between the V1 capillary RBC flux and PO_2_ ratio. The linear regression slope = 0.01 s·RBC^-1^ (R^2^≈0.12).

The RBC flux in the A1-A3 capillaries (97±6 RBC/s) was ∼2.7 times higher than that in the V1-V3 capillaries (36±2 RBC/s; Fig. 7c). Since blood flow obeys the principle of mass conservation (i.e. the numbers of RBCs entering the capillary paths per unit time on the arteriolar side and exiting from the venous side must be equal), this result is in agreement with the previously observed ∼3-fold-greater number of V1 than A1 capillaries in mouse cortex (Nguyen et al., 2011). Furthermore, the RBC speed in the A1-A3 capillaries (1.9±0.2 mm/s) had a similar ratio (∼3.2 times) to that in the V1-V3 capillaries (0.6±0.1 mm/s; Fig. 7e), which is also consistent with the strong correlation between RBC flux and speed (Supplementary Fig. 6) (Desjardins et al., 2014; Kleinfeld et al., 1998; Lyons et al., 2016).

We found strong positive correlations between the capillary RBC flux and both Mean-PO_2_ and SO_2_ in the downstream (V1-V3) capillaries, but not in the upstream (A1-A3) capillaries (Fig. 7g, h), suggesting that a positive correlation between the RBC flux and oxygenation may be gradually building up along the capillary paths. In addition, we observed very heterogeneous distributions of both Mean-PO_2_ (from ∼11 mmHg to ∼68 mmHg) and RBC flux (from ∼30 RBCs/s to ∼110 RBCs/s) in the V1 capillaries (Fig. 7i, j). Lastly, the V1 capillary RBC flux was correlated positively with the ratio of the V1 capillary Mean-PO_2_ to the PO_2_ in the adjacent post-capillary venules (PCV PO_2_) (Fig. 7k). Therefore, while heterogeneous oxygen delivery is taking place along different capillary paths, the capillary paths with higher RBC flux are oxygenated, contributing more to the increase in the PCV oxygenation.

Finally, EATs observed in the A1-A3 capillaries (10.9±1.9 mmHg) were noticeably smaller than in the V1-V3 capillaries (16.6±2.3 mmHg; Fig. 7c), although this difference did not reach statistical significance (P = 0.09). In both this work (Supplementary Fig. 5) and a previous study (Lyons et al., 2016), EATs did not exhibit any obvious dependence on the Mean-PO_2_, RBC flux or speed, but they were correlated negatively with hematocrit. Nevertheless, the observed trend that the average EATs amplitude in the A1-A3 capillaries was smaller than that in the V1-V3 capillaries is unlikely to be related to hematocrit, as we did not observe significant difference between the upstream and downstream hematocrit values (Fig. 7f).

## DISCUSSION

We have performed measurements of absolute intra-vascular PO_2_ in arterioles, venules and a large number of capillaries (6,544 capillaries across n = 15 mice), as well as EATs, RBC flux, speed and hematocrit in a subset of capillaries (978 capillaries across n = 15 mice; Supplementary Fig. 2). These parameters were measured simultaneously in the whisker barrel cortex in head-restrained awake C57BL/6 mice and thus were free of the confounding effects of anesthesia on neuronal activity, CBF and brain metabolism.

A new two-photon-excitable phosphorescent oxygen probe was used in this study. This probe belongs to the class of dendritically protected oxygen probes (Lebedev et al., 2009), and it is built around a newly developed Pt tetraarylphthalimidoporphyrin (PtTAPIP) (Esipova et al., 2017). In comparison to its predecessor (PtP-C343) (Finikova et al., 2008), the new probe exhibits red-shifted two-photon excitation and emission maxima, higher phosphorescence quantum yield and larger two-photon absorption crosssection. Furthermore, in contrast to PtP-C343 the phosphorescence decay of the new probe exhibits well-defined single-exponential kinetics, which aids to improve accuracy of oxygen quantification. The above properties of the probe facilitated large-scale sampling of both capillary RBC flow and PO_2_ deeper in the brain, e.g., 600 μm vs. 450 μm (Lyons et al., 2016; Sakadžić et al., 2014), and with higher signal-to-noise ratios, which, in turn, powered the statistical analyses beyond what was achievable with previous probes (Devor et al., 2011; Lecoq et al., 2011; Lyons et al., 2016; Parpaleix et al., 2013; Sakadžić et al., 2010, 2014). In addition, the very high phosphorescence quantum yield of PtTAPIP (Esipova et al., 2017) enabled detection of RBC passages in capillaries based on phosphorescence alone, without the need to enhance the signal from the blood plasma using additional chromophores (Lyons et al., 2016).

We first assessed oxygenation in the diving arterioles and surfacing venules and observed a decrease in PO_2_ with cortical depth for both vessel types (Fig. 2a). The PO_2_ decrease in the diving arterioles between layer I and V was about 15 mmHg, equivalent to the difference in SO_2_ of 3.8 %. This modest oxygen extraction along the diving arterioles implies that much more oxygen was extracted from the rest of the microvascular network, including arteriolar branches, capillaries and potentially venules. Previous measurements performed in mice (Kisler et al., 2017; Moeini et al., 2018) and rats (Devor et al., 2011; Sakadžić et al., 2016) reported pronounced PO_2_ gradients in the periarteriolar tissue around the cortical diving arterioles, implying extraction of oxygen from cortical diving arterioles. Specifically, a significantly steeper decrease in PO_2_ with cortical depth was observed directly in diving arterioles in isoflurane-anesthetized mice by Kazmi et al. (Kazmi et al., 2013) and indirectly, based on the depth dependence of the extravascular (tissue) PO_2_ measured immediately next to the diving arterioles, in α-chloralose anesthetized rats by Devor et al. (Devor et al., 2011) The most likely reason for these discrepancies is the higher suppression of CBF than CMRO_2_ under different anesthesia regimes. In awake mice, Lyons et al. (Lyons et al., 2016) measured approximately constant PO_2_ in the diving arterioles over cortical depth of 0-400 μm. This result is in agreement with our current observation and implies a small loss of O_2_ along the diving arterioles in awake mice.

In contrast to the moderate PO_2_ decrease along the penetrating arterioles, the decrease in the venular PO_2_ from layer I to V was much smaller (∼6 mmHg), but resulted in a larger fractional decrease in SO_2_ (8.3 %) than in the penetrating arterioles (Fig. 2b). An increase in PO_2_ in ascending venues towards the cortical surface has been previously observed in isoflurane-anesthetized mice (Sakadžić et al., 2010). However, the PO_2_ values, measured previously along the ascending venules in the fore-paw and hind-paw regions of the somatosensory cortex in awake mice, were reported to be nearly constant (Lyons et al., 2016). The difference between these observations may be attributed to the difference in the studied cortical regions and/or to the lower measurement accuracy associated with the previous oxygen probe because of its lower brightness and inability to resolve small PO_2_ changes. In addition, some discrepancies in the absolute baseline PO_2_ in arterioles and venules at different depths were observed between our present and some previous studies (Kazmi et al., 2013; Lecoq et al., 2011; Lyons et al., 2016; Parpaleix et al., 2013; Sakadžić et al., 2010). Different anesthesia regimes, different cortical areas and, most importantly, use of anesthetized vs. awake animals would be the most obvious factors underlying the observed discrepancies. Furthermore, cortical temperature typically was not controlled in the previous studies, while the presence of a cranial window can cause reduction in the cortical temperature (Shirey et al., 2015), especially when imaging is carried out using a non-heated water-immersion objective. Temperature affects all physiological parameters and chemical properties, including O_2_ diffusion and solubility, hemoglobin affinity to O_2_, CBF, CMRO_2_ and the triplet decay time of the oxygen probe (Benesch et al., 1969; Croughwell et al., 1992; Finikova et al., 2008; Hanks and Wallace, 1949; Jones and Siegel, 1969; PRAY, 1952; Rosomoff and Holaday, 1954; Rossing and Cain, 1966; Soukup et al., 2002) – all of which may affect the experimental results.

A faster decrease in SO_2_ with cortical depth in the ascending venules than in the penetrating arterioles resulted in a higher depth-dependent OEF in the deeper cortical layers, reaching the maximum in layer IV (Fig. 2b). This implied that oxygen extraction was higher in deeper cortical layers, which would be in agreement with the finding that layer IV has the highest neuronal and capillary density in mouse cortex (Blinder et al., 2013; Lefort et al., 2009; Patel, 1983; Wu et al., 2016), and that the cells in layer IV exhibit the highest cytochrome oxidase labeling activity, suggestive of the highest oxidative metabolism (Land and Simons).

The mean capillary RBC flux and speed decreased only slightly with cortical depth, without reaching statistically significant difference across layers I-V (Fig. 3a, d). This might be due to the redistribution of the blood flow over the denser capillary network in the deeper cortical layers, especially in layer IV (Blinder et al., 2013; Sakadžić et al., 2014; Wu et al., 2016), causing the RBC flux and speed to be lower in individual capillaries. The observed mean capillary RBC flux (41.5±1.2 RBCs/s; Supplementary Fig. 2) is in good agreement with the value of 41.9±1.8 RBCs/s, reported by Lyons et al. (Lyons et al., 2016), and somewhat lower than ∼48 RBC/s reported by Moeini et al. (Moeini et al., 2018) for somatosensory cortex in awake mice. The average capillary Mean-PO_2_ increased slightly by ∼7 mmHg from layer I to V (Fig. 3g), which is in agreement with the previously reported increase in capillary Mean-PO_2_ in both awake (Lyons et al., 2016) and isoflurane-anesthetized mice (Sakadžić et al., 2014) across the upper 400 μm of the cortex. One potential explanation for this increase could be related to the finding by Blinder et al. (Blinder et al., 2013), who reported that, in the upper 600 μm of somatosensory cortex, the probability of penetrating arterioles giving off side branches increased with cortical depth, whereas the probability for the ascending venules decreased with depth. This suggests that the ratio between the number of the more oxygenated (upstream) capillary branches and that of the less oxygenated (downstream) capillary branches increases with depth, contributing to the higher capillary PO_2_ at greater depth.

Importantly, we found that the distributions of capillary RBC flux, speed, and Mean-PO_2_ undergo homogenization in cortical layers IV and V (Fig. 3). Both analytical and more accurate numerical models of oxygen advection and diffusion in microvascular networks predict that capillary blood flow homogenization facilitates oxygen delivery to tissue (Jespersen and Østergaard, 2012; Østergaard et al., 2013, 2014a, 2014b; Rasmussen et al., 2015). Indeed, homogenization of capillary blood flow, measured as capillary transit time heterogeneity and STD of capillary RBC flux during functional brain activation in healthy anesthetized animals, have been reported previously (Gutiérrez-Jiménez et al., 2016; Lee et al., 2015; Stefanovic et al., 2007). Interestingly, capillary flow homogenization was absent in the mouse model of Alzheimer’s disease (Gutiérrez-Jiménez et al., 2018), while increased capillary blood flow heterogeneity was found in patients of Alzheimer’s disease (Eskildsen et al., 2017; Nielsen et al., 2017), suggesting a possible link between disturbed capillary flow patterns, reduced oxygen supply to tissue, and the progression of neurodegeneration. Our observations are in agreement with the previously reported lower capillary transit time heterogeneity in cortical layers IV and V than in the upper layers in anesthetized mice (Merkle and Srinivasan, 2016). Furthermore, capillary Mean-PO_2_ also exhibited a similar homogenization trend (Fig. 3h, i). This could be expected since capillary RBC flow is correlated positively with Mean-PO_2_ (Lyons et al., 2016).

Temporal fluctuations of the resting state capillary RBC flux, speed, and Mean-PO_2_, measured during 9-s-long periods, were found to be attenuated in the deeper cortical layers (Fig. 4). The temporal resolution of our measurements (0.6 s per single point) ensured that the blood flow and oxygenation fluctuations due to cardiac and respiratory cycles, as they occurred in awake mice, were averaged out, but it was sufficient to capture the dynamics related to the rate of oxygen consumption, which could be estimated as the time for tissue PO_2_ to drop to zero after blood flow stoppage. As reported in cats, that time was at least several seconds (Acker and Lübbers, 1977; Whalen and Nair, 1975). The cause of the dampening of the temporal fluctuations with depth is unclear. It may be related to the anatomical and functional differences between cortical layers; it also can be due to the expected dampening along the vascular paths if the temporal fluctuations originate from the upstream arteriolar blood flow.

We observed strong positive correlations between the temporal fluctuations of capillary RBC flux and Mean-PO_2_ (Fig. 5). This observation is in agreement with the previously reported positive correlation between the fluctuations of RBC flux and extravascular (tissue) PO_2_ next to the capillaries in cancerous tumors in rats, although the latter occurred at a lower frequency range (Braun et al., 1999; Kimura et al., 1996). Importantly, the fluctuation of capillary Mean-PO_2_ was found to be correlated positively with the fluctuation of RBC flux significantly stronger than with the fluctuations of RBC speed or hematocrit (Fig. 5b, c). This result could be expected, since most oxygen in the blood is bound to the hemoglobin inside RBCs (Pittman, 2011). We also observed strong positive correlation between the fluctuations of RBC flux and speed, but the fluctuation of hematocrit was poorly correlated with both of them (Supplementary Fig. 4). These correlations between the temporal fluctuations are in agreement with the trends measured using the populations of capillaries (Supplementary Fig. 6) as well as with the previous studies (Kleinfeld et al., 1998; Santisakultarm et al., 2012; VanTeeffelen et al.). Despite poor correlation between the RBC flux and hematocrit, we observed reasonably strong positive correlation between the temporal fluctuations of capillary Mean-PO_2_ and hematocrit (Fig. 5). Similarly, capillary Mean-PO_2_ and hematocrit, measured in the populations of capillaries, were also positively correlated shown in Supplementary Fig. 6 and in (Lyons et al., 2016) emphasizing the specific role of hematocrit in oxygen transport (Lücker et al., 2017).

Longitudinal capillary PO_2_ gradients were measured to assess the intracapillary resistance to oxygen delivery to tissue and to calculate the capillary SO_2_ (Fig. 6). For measuring EATs, we followed the previously developed methods (Lecoq et al., 2011; Lyons et al., 2016; Parpaleix et al., 2013). Compared to the previous studies performed with the old probe PtP-C343 (Lecoq et al., 2011; Lyons et al., 2016; Parpaleix et al., 2013), the higher brightness of the new probe enabled us to measure PO_2_ gradients in a much larger number of capillaries (∼1,000) over a greater cortical depth (down to 600 μm below the cortical surface) and using a shorter acquisition time (9 s) per measurement location. In agreement with Lyons et al. (Lyons et al., 2016), we observed EATs in capillaries (Fig. 6), although the magnitude was lower than previously reported. EATs did not appear to depend significantly on the RBC flux, speed or Mean-PO_2_, but were correlated negatively with the hematocrit (Supplementary Fig. 5). Our measurements are in agreement with the previous observations (Lyons et al., 2016) as well as with the theoretical predictions (Hellums, 1977; Lücker et al., 2017). Importantly, we detected a significant decrease in EATs towards deeper cortical layers, dropping from 13.4 mmHg in cortical layer I to 9.9-11.0 mmHg in layers IV and V. Since EATs are directly related to the intravascular resistance to the diffusive oxygen transport from RBCs to tissue (Hellums, 1977), reduced EATs amplitude in the deeper layers represents another example of the adaptation of the microvascular network in order to facilitate local oxygen delivery in cortical regions with higher oxygen demand (Pries et al., 1994, 1998; Reglin et al., 2009, 2017).

The elevated SO_2_ in the surfacing venules in layers I-III suggests that in the upper cortical layers (e.g. layers I-III) the downstream capillaries and post-capillary venules that feed the surfacing venules are more oxygenated compared to the surfacing venules in the deeper layers (e.g. layers IV and V). We therefore assigned branching orders to the capillary segments in cortical layers I-III and investigated the distributions of capillary RBC flow and oxygenation along the capillary paths. We found that the average SO_2_ in the V1 capillaries selected across layers I-III was indeed higher than the SO_2_ in the ascending venules in both layers I and II/III, confirming that high oxygen content in the superficial capillaries contributed to the increase in the venular SO_2_ towards brain surface. This finding also suggests that the baseline oxygen extraction may be different between the cortical layers. If true, this may have implications on the interpretation of results from different imaging modalities such as BOLD fMRI (Blockley et al., 2015; Griffeth and Buxton, 2011; Siero et al., 2011; Silva and Koretsky, 2002; Vazquez et al., 2006; Yu et al., 2014). Secondly, approximately 1/3 of the total oxygen extraction took place along the paths from the pial arterioles to the A1 capillary segments in layers I-III. This observation differs from our previous study in isoflurane-anesthetized mice, where we found, based on the measurements in the upper 450 μm of somatosensory cortex, that ∼50 % of the oxygen delivered to brain tissue was extracted from arterioles (Sakadžić et al., 2014). The discrepancy might be due to the effect of anesthesia, which suppressed cerebral oxygen metabolism and possibly increased both CBF and arteriolar surface area due to vasodilation (Alkire et al., 1999; Goldberg et al., 1966; Ogawa et al., 1990). As a result of these perturbations, oxygen extraction could be shifted towards the upstream microvascular segments (Sakadžić et al., 2014). Based on the observed difference in oxygen extraction and the assumed difference in oxygen metabolism between cortical layers, we anticipate that in cortical layers IV and V, the fraction of the extracted oxygen from arterioles may be smaller than in layers I-III, but still significant.

Our data provided additional evidence that the distributions of capillary flow and oxygenation were highly heterogeneous and strongly positively correlated with one-another, in particular in the downstream capillaries (Fig. 7). This implies that the mixed venous blood oxygenation was, therefore, a result of the contributions from the wide distribution of capillary paths, carrying blood that was both more and less oxygenated than the blood in the postcapillary venules (Fig. 7). At the extreme ends of this distribution are capillary paths with very low and very high oxygenation and blood flow. The paths with very low oxygenation may be the first sites of coupling of microvascular dysfunction and progression of various brain pathologies (Erdener et al., 2017; Lücker et al., 2018). Indeed, it has been recently reported that only ∼0.4 % of the cortical capillaries in healthy mice had ‘stalled flow’, but in transgenic mice of Alzheimer’s disease the fraction was as high as 2 % (Hernandez et al., 2016; Momjian-Mayor and Baron, 2005). As an example of such poorly oxygenated/perfused capillaries, we identified a capillary segment with PO_2_ of ∼15 mmHg and no detectable RBC flow during the 9-s-long acquisition (Supplementary Fig. 7). In contrast, the paths with very high oxygenation and blood flow (Supplementary Fig. 8) may be especially good sites for implementing blood flow control in response to locally increased oxygen demand, since increasing their resistance to flow may quickly redistribute blood over the nearby capillary network. However, the prevalence of such highly oxygenated/perfused capillaries in the cortical microvascular network and the existence of mechanisms of their site-specific control are unknown (Hudetz et al., 1996). Further studies should address these questions in relation to both normal and pathological brain conditions.

One potential limitation of this study is that the assignment of the cortical layers as a function of the cortical depth was performed based on the literature data (Blinder et al., 2013; Lefort et al., 2009), as opposed to identification of layer-specific anatomical landmarks. Nevertheless, while slight shifts in the layer boundaries, which may have resulted from such an assignment, may have affected the exact values, it is unlikely that the observed general trends and significant differences in the observables between the layers would have changed. In addition, layer V in mouse barrel cortex spans the depth range of 450-700 μm (Blinder et al., 2013; Lefort et al., 2009), so that the range of 450-600 μm, interrogated in this study, likely overlaps the best with the upper part of layer V (i.e. layer V_a_) (Blinder et al., 2013; Lefort et al., 2009). Another limitation is that capillaries as a vessel type and their branching order indices were identified based on their diameter, without taking into account the smooth muscle cell coverage and pericyte types (Attwell et al., 2016; Hall et al., 2014; Peppiatt et al., 2006). In principle, misclassifying vessel types and/or branching order indices could influence our analysis. However, based on the overwhelmingly larger number of capillaries compared to the non-penetrating arterioles and venules, our conclusions in general are unlikely to be different. Furthermore, identification of capillaries based on the biochemical staining of smooth muscle cells and pericytes (Hall et al., 2014; Hartmann et al., 2015) will add significant complexity to the study, especially considering a large sample size used in our analysis, but in the end may not provide more accurate results, as the debate on what is a proper classification of vessel-types based on staining is still ongoing (Hill et al., 2015).

In conclusion, we have experimentally mapped the distributions of microvascular flow and oxygenation in the whisker barrel cortex in awake mice using two-photon phosphorescence lifetime microscopy. We have found evidence that oxygen was extracted differently in different cortical layers, and that the distributions of capillary blood flow and oxygenation were adjusted across layers in a way that facilitated oxygen delivery in the deeper layers (Table 1). Specifically, the depth-dependent OEF was measured higher in cortical layers IV and V, where the oxidative metabolism is presumably the highest in cortex. This increase was accompanied by the homogenization of capillary RBC flow and oxygenation, as well as by the reduction of intracapillary resistance to oxygen diffusion to tissue (inferred from the changes of EATs amplitude). In addition, we have found that arterioles in the superficial cortical layers (e.g. layers I-III) contributed significantly to the oxygen extraction from blood (34 %) even in awake mice. We anticipate that our results and analysis will help better understand the normal brain physiology, and the progression of brain pathologies that affect cerebral microcirculation. They will also inform more accurate biophysical models of the cortical layer-specific oxygen delivery and consumption, as well as improve the interpretation of the results from other brain imaging modalities.

**Table 1.** Summary of microvascular oxygenation and RBC flux changes with cortical depth. The arrows pointing up (↑) and down (↓) indicate increase and decrease of the measured observables from cortical layers I-III to layers IV/V, respectively. The fields marked in blue represent the parameters reported in the literature. The fields marked in black represent our experimental measurements.

| *Parameter* | *Change with depth* |
| --- | --- |
| **Neuronal density** (Blinder et al., 2013; Lefort et al., 2009; Wu et al., 2016) | ↑ |
| **Capillary density** (Blinder et al., 2013; Wu et al., 2016) | ↑ |
| **Depth-dependent OEF** | ↑ |
| **Capillary RBC flux STD/CV** | ↓ |
| **Capillary PO<sub>2</sub> STD/CV** | ↓ |
| **EATs** | ↓ |

## MATERIALS AND METHODS

### Animal preparation

We used n = 15 C57BL/6 mice (3-5 months old, 20-25 g, female, Charles River Laboratories) in this study. We followed the procedures of chronic cranial window preparation outlined by Goldey *et al.* (Goldey et al., 2014) A custom-made head-post (Mateo et al., 2011) allowing repeated head immobilization was glued to the skull, overlaying the right hemisphere (Supplementary Fig. 1). A craniotomy (round shape, 3 mm in diameter) was performed over the left hemisphere, centered approximately over the E1 whisker barrel. The dura was kept intact. The cranial window was subsequently sealed with a glass plug (Komiyama et al., 2010) and dental acrylic. After surgery, mice were given 5 days to recover before starting the habituation training. Mice were gradually habituated to longer periods (from 10 minutes to 2 hours) of head-restraint, resting on a suspended bed made of soft fabric, in a home built platform, under the microscope. They were rewarded with sweetened milk every 15 minutes during both training and experiments. While head-restrained, the mice were free to readjust their body position and from time to time displayed natural grooming behavior. All animal surgical and experimental procedures were conducted following the Guide for the Care and Use of Laboratory Animals and approved by the Massachusetts General Hospital Subcommittee on Research Animal Care (Protocol No.: 2007N000050).

### Two-photon microscope

In this study, we employed our previously developed home-built two-photon microscope (Fig. 1a) (Sakadžić et al., 2010; Yaseen et al., 2015). Briefly, a pulsed laser (InSight DeepSee, Spectra-Physics, tuning range: 680 nm to 1300 nm, ∼120 fs pulse width, 80 MHz repetition rate) was used as an excitation source. Laser power was controlled by an electro-optic modulator (EOM). The laser beam was focused with a water-immersion objective lens (XLUMPLFLN20XW, Olympus, NA = 1.0), and scanned in the X-Y plane by a pair of galvanometer scanners. The objective was moved along the Z axis by a motorized stage (M-112.1DG, Physik Instrumente) for probing different cortical depth. The phosphorescent oxygen-sensitive probe was excited at 950 nm. The emitted phosphorescence, centered at 760 nm, was directed towards a photon-counting photomultiplier tube (H10770PA-50, Hamamatsu) by a dichroic mirror (FF875-Di01-25×36, Semrock), followed by an infrared blocker (FF01-890/SP-25, Semrock) and an emission filter (FF01-795/150-25, Semrock). During the experiments, the objective lens was heated by an electric heater (TC-HLS-05, Bioscience Tools) to maintain the temperature of the water between the cranial window and the objective lens at 36-37 °C. In addition, we attached the sensor of an accelerometer module to the suspended bed of the mouse platform to record the signals induced by mouse motion. Mice were continuously monitored during experiments by acquiring live videos with a CCD camera (CoolSNAPfx, Roper Scientific) using a LED illumination at 940 nm.

### Intravascular PO_2_ imaging

Before imaging, mice were briefly anesthetized by isoflurane (1.5-2 %, during ∼2 minutes), and then the solution of the phosphorescent oxygen probe (0.1 ml at ∼34 μM) was retro-orbitally injected into the bloodstream. Mice were recovered from anesthesia and then head fixed under the microscope. The imaging session was started 30-60 minutes after the probe injection and lasted for up to 2 hours.

The acquisition protocol is illustrated in Fig. 1b. We first recorded a CCD image of the mouse brain surface vasculature (Fig. 1d) under green light illumination to guide the selection of a region of interest for the subsequent functional imaging. The two-photon microscopic measurements were collected at different X-Y planes, perpendicular to the optical axis (Z). The imaging planes were separated in depth by ≤50 μm, spanning the depth range from cortical surface to 600 μm under cortical surface. At each imaging depth, we first performed a raster scan of phosphorescence intensity, which revealed the locations of the micro-vessels within a 500 × 500 μm^2^ field of view (FOV). Then, we manually selected the measurement locations inside all the vascular segments captured within the FOV. At each selected location, the phosphorescent oxygen probe in the focal volume was excited with a 10-μs-long laser excitation at 950 nm gated by the EOM, followed by a 290-μs-long collection of the emitted phosphorescence. Typically, at each location, such 300-μs-long excitation/decay cycle was repeated 2,000 times (0.6 s) to obtain an average phosphorescence decay with sufficient signal-to-noise ratio (SNR) for an accurate lifetime calculation.

### Imaging of EATs and capillary RBC flow

In a subset of capillaries, we repeated the 300-μs-long excitation/decay cycle 30,000 times at each measurement location, corresponding to a 9-s-long acquisition per capillary (Lecoq et al., 2011). This long acquisition time allowed us to estimate EATs by grouping the phosphorescence decays as a function of their distance to their nearest RBC center, while ensuring a sufficient number of phosphorescence decays associated with each distance bin for averaging. The same dataset was used to calculate capillary RBC flux, as well as to estimate RBC speed and hematocrit averaged over the 9-s-long intervals. In addition, the same dataset was used to estimate temporal fluctuations of capillary RBC flux, speed, hematocrit, and capillary PO_2_ during the 9-s-long periods.

### Acquisition of microvascular angiograms

Microvascular angiograms were acquired by two-photon microscopic imaging of blood plasma labeled with dextran-conjugated Sulforhodamine-B (SRB) (0.15-0.2 ml at 5 % W/V in saline, R9379, Sigma Aldrich). The SRB solution was retro-orbitally injected into the blood stream under brief isoflurane anesthesia (1.5-2 %, during ∼2 minutes). The microvascular stacks were acquired within a 700 × 700 μm^2^ FOV, centered over the same region of interest as for PO_2_ imaging. In each mouse, the angiogram and intravascular PO_2_ were acquired separately on different days in order not to stress the animals by the combined long imaging session.

### Calculation of PO_2_

We rejected the initial 5-μs phosphorescence decay data after the 10-μs-long excitation gate, and used the remaining 285-μs decay to fit for the phosphorescence lifetime. The phosphorescence lifetime was calculated by fitting the average phosphorescence decay to a single-exponential decay function, using a standard non-linear least square minimization algorithm (Finikova et al., 2008; Sakadžić et al., 2010). The lifetime was converted to absolute PO_2_ using a Stern-Volmer type calibration plot obtained in an independent oxygen titration experiment, conducted with the same 300-μs-long excitation/decay acquisition protocol as in our *in vivo* recordings.

### Calculation of capillary RBC flux, speed, and hematocrit

The phosphorescence intensity was calculated by integrating the phosphorescence photon counts over each 300-μs-long excitation/decay cycle. Since the phosphorescent probe is confined to blood plasma, but not permeates RBCs, as all the dendritic oxygen probes (Lebedev et al., 2009), the variations of the phosphorescence intensity recorded in capillaries encoded the passing of RBCs through the optical focus (Lecoq et al., 2011; Parpaleix et al., 2013) (Fig. 1b, c). Following the previously described procedures (Lecoq et al., 2011; Parpaleix et al., 2013), the phosphorescence intensity time course was segmented using a binary threshold method. A representative phosphorescence intensity time course is shown in Fig. 1c, where a RBC and a blood-plasma-passage induced phosphorescence intensity transients are denoted by arrows. Subsequently, capillary RBC flux was calculated by counting the number of detected RBCs (i.e. valleys in the binary segmented curve) during the acquisition time. Hematocrit was estimated as the ratio of the combined duration of all valleys associated with the RBC passages to the duration of the entire time course. Finally, RBC speed for each RBC passage event was estimated as v = ø/Δt, where Δt is the time for the RBC to pass through the focal zone, and 0 is RBC diameter, assumed to be 6 μm (Unekawa et al., 2010). For each capillary, the mean RBC speed was calculated by averaging over all RBC passage events throughout the time course.

### Calculation of capillary PO_2_ gradients and EATs

The phosphorescence decays from the 9-s-long acquisition in each capillary were grouped by their distance to the nearest RBC center using 1-μm-wide bins. The distance to the nearest RBC center was estimated as v·Δt′, where v was the speed of the nearest RBC, and Δt′ was the time interval between the phosphorescence decay event and the center of the nearest RBC passage (i.e. the center of the valley in the time course). The decays at each 1-μm-wide bin were subsequently averaged, and then PO_2_ calculated using the procedure described above.

The phosphorescence decays at the RBC passages (i.e. valleys in the binary segmented time course) and in proximity to the mid-points between adjacent RBCs (corresponding to the central 40 % of the peaks in the binary segmented time course) were averaged to calculate RBC-PO_2_ and InterRBC-PO_2_, respectively. The erythrocyte-associated transients of capillary PO_2_ (EATs) were calculated as EATs = RBC-PO_2_ – InterRBC-PO_2_. Mean-PO_2_ was calculated by averaging all the phosphorescence decays acquired within a capillary, regardless of their distance to RBCs (Lecoq et al., 2011; Parpaleix et al., 2013; Lyons et al., 2016).

### Calculation of SO_2_ and depth-dependent OEF

The oxygen saturation of hemoglobin (SO_2_) was computed based on PO_2_ using the Hill equation with the parameters (h = 2.59, P_50_ = 40.2 mmHg) specific for C57BL/6 mice (Uchida et al., 1998). Here, h is the Hill coefficient, and P50 is the oxygen tension at which hemoglobin is 50 % saturated. SO_2_ in the penetrating arterioles and surfacing venules was calculated based on their Mean-PO_2_. SO_2_ in capillaries was calculated based on the RBC-PO_2_ (Lyons et al., 2016; Sakadžić et al., 2014).

The depth-dependent OEF (DOEF), in a given cortical layer, was calculated as (SO_2,A_–SO_2,V_)/SO_2,A_, where SO_2,A_ and SO_2,V_ represent the layer-specific SO_2_ in the diving arterioles and surfacing venules, respectively. Therefore, DOEF in a certain layer measures the OEF accumulated downstream from that layer and, as a special case, DOEF in layer I represents the global OEF in the interrogated cortical tissue territory.

### Quantification of the temporal fluctuations of capillary Mean-PO_2_, RBC flux, speed, and hematocrit

The phosphorescence decays recorded during the 9-s-long acquisition (30,000 repetitions of the excitation/decay cycle) in each capillary were divided into 15 groups using 0.6-s-long bins (2,000 repetitions for each bin). RBC flux, speed, hematocrit, and Mean-PO_2_ were calculated with the 2,000 phosphorescence decays at each bin, yielding 15-point time courses of these 4 parameters for each assessed capillary. Subsequently, for each of the 4 parameters, we quantified the temporal fluctuation by computing the standard deviation (STD) and coefficient of variance (CV) from the 15-point data. Here, CV is defined as the ratio of STD to mean (Golub and Pittman, 2005).

### Identification of capillary branching order

Capillaries were identified empirically based on morphology and vascular diameters (Cai et al., 2018; Sakadžić et al., 2014). Their branching order indices were assigned by visually inspecting the microvascular angiograms. Starting immediately after the pre-capillary arteriole (PCA), capillary segments were counted in the direction of blood flow and indexed as Ai (i = 1, 2, 3, …). Analogously, starting immediately before the post-capillary venule (PCV), capillary segments were counted in the opposite direction of blood flow and indexed as Vi (i = 1, 2, 3, …). In the analysis, only the first 3 upstream capillary segments (A1-A3) and the last 3 downstream capillary segments (V1-V3) were considered, as visually inspecting and confirming higher branching orders of capillaries would be more challenging. In addition, most of the capillaries selected were within the central part of the FOV of the microvascular angiograms and in the cortical depth of ≤300 μm, which were due to the difficulty in tracking the capillaries close to the boundaries of the FOV and at greater depth. PCAs and PCVs were identified based on their morphology, vascular diameters, and PO_2_ values, and confirmed by tracing the vessels from the brain surface with the microvascular angiograms (Sakadžić et al., 2014).

### Cortical layer-specific data analysis

The capillary RBC flow and PO_2_ properties acquired in each animal were grouped into 4 groups based on the cortical depth: 0-100 μm, 100-320 μm, 320-450 μm, and 450-600 μm. These depth ranges approximately correspond to the cortical layers I, II/III, IV, and V, respectively, in the whisker barrel cortex in 3-month-old C57BL/6 mice (Blinder et al., 2013; Lefort et al., 2009). For each cortical layer in each mouse, the absolute values of capillary RBC flow and PO_2_ properties were averaged, and the STD and CV computed. Subsequently, the measurements belonging to each cortical layer were averaged over mice.

### Rejection of motion artefacts

Data affected by mouse motion were rejected based on the signal generated by the accelerometer attached to the fabric underneath the mouse. We excluded from analysis the phosphorescence decays acquired within the time intervals determined by an empirically defined threshold of the accelerometer signal amplitude. In addition, visual inspection of the phosphorescence intensity traces acquired in the microvascular segments was also used to find and reject motion artefacts. This was achieved by looking for the sudden changes of phosphorescence intensity or loss of contrast between RBC and plasma passing through the focal volume. Finally, long episodes of motion were captured by the live-videos recorded by a CCD camera during acquisition. When motion occurred, the acquisition was manually stopped and the corresponding measurements were excluded from the analysis.

### Construction of the composite image

To construct composite images such as the one shown in Fig. 1g, tubeness filtering (Sato et al., 1998) and intensity thresholding were applied to segment the two-photon angiograms into binary images. Subsequently, PO_2_ measurements were spatially co-registered with the segmented angiogram. The three-dimensional composite image was created by color-coding the experimental PO_2_ values in the corresponding vascular segments (shades of gray). The color-coding was performed by assigning the PO_2_ (or Mean-PO_2_ for capillary) value measured in the focal volume within a vascular segment to the whole segment. For some vascular segments without PO_2_ measurements, such measurements were instead available for the segments joining them in each end. To such segments, we assigned the average PO_2_ values of the connecting segments.

### Statistical analysis

Statistical comparisons were made using Student’s t-test (MATLAB, MathWorks Inc.). P value less than 0.05 was considered statistically significant. Specifically, in Figs. 3, 4, and 6c, pairwise t-test was performed between the parameters in layer I and those in each of other layers; in Fig. 5c, pairwise t-test was performed between the flux-PO_2_ correlation and each of the other correlations. Further details about the statistical information of measurements are provided in the text and figure legends, where relevant.

Mean values and standard deviations of parameters needed to estimate the sample size were either assumed to be the same as measured previously in anesthetized animals or assumed empirically. Since multiple measurements were performed in the same animals, sample size (i.e., n=15 mice) was set based on anticipation that the most demanding one will be to detect 30% difference between mean EATs values (coefficient of variance = 0.3, power = 0.8, α=0.05).

## DATA AVAILABILITY

All data generated or analyzed in support the findings of this study are within this paper.

## ACKNOWLEDGEMENTS

Support of the grants R01NS091230, MH111359, EB018464, NS092986 and P01NS055104 from the National Institutes of Health, USA, is gratefully acknowledged.

## AUTHOR CONTRIBUTIONS

**Conceptualization:** B.L., S.S. and D.A.B.; **Data curation:**B.L. and S.S.; **Formal analysis:** B.L., S.S. and A.D.; **Funding acquisition:** S.S., A.D., D.A.B. and S.A.V.; **Investigation:** B.L. and S.S.; **Methodology:** S.S., S.A.V., D.A.B., A.D., T.V.E., M.A.Y., B.L., K.K., B.F., M.M. and I.S.; **Project administration:** S.S.; **Resources:** S.S.; **Software:** S.S., B.L., M.A.Y. and S.K.; **Supervision:** S.S.; **Validation:** S.A.V., L.Ø., A.D., D.A.B., F.L., M.D. and M.M.; **Visualization:** S.S., B.L. and S.K.; **Writing—original draft:** B.L.; **Writing—review and editing:** S.S., B.L., S.A.V., A.D., L.Ø., D.A.B., F.L., M.D. and M.M.

## SUPPLEMENTARY INFORMATION

Supplementary results are available for this paper.

## SUPPLEMENTARY RESULTS

**Supplementary Figure 1.**
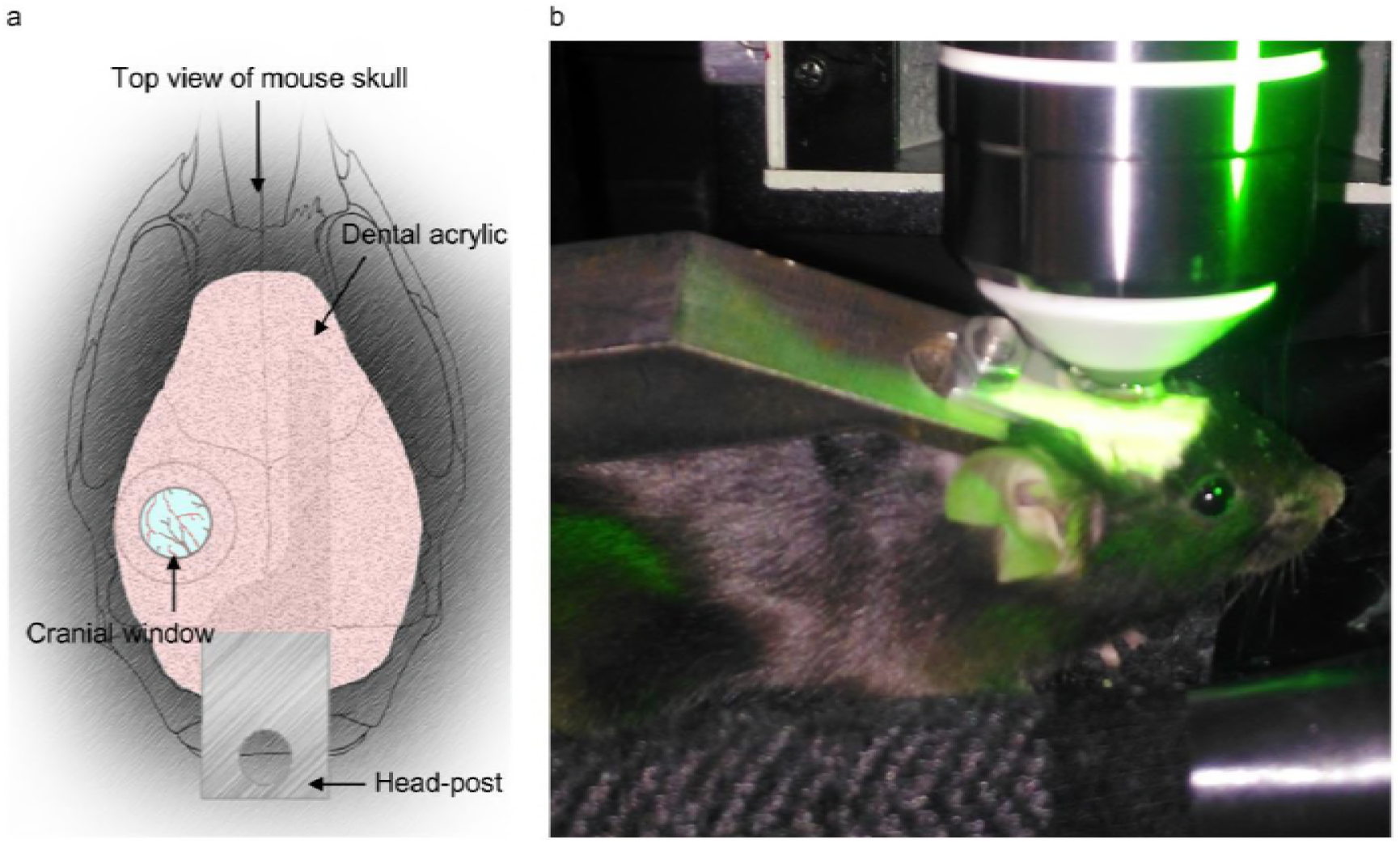
Schematic diagram of a chronic cranial window and photograph of an awake head-restrained mouse. **a.** Schematic diagram of a chronic cranial window on mouse skull. **b.** Photograph of an awake head-restrained mouse on the training platform, snapshotted prior to imaging experiment using a CCD camera, under the two-photon microscope.

**Supplementary Figure 2.**
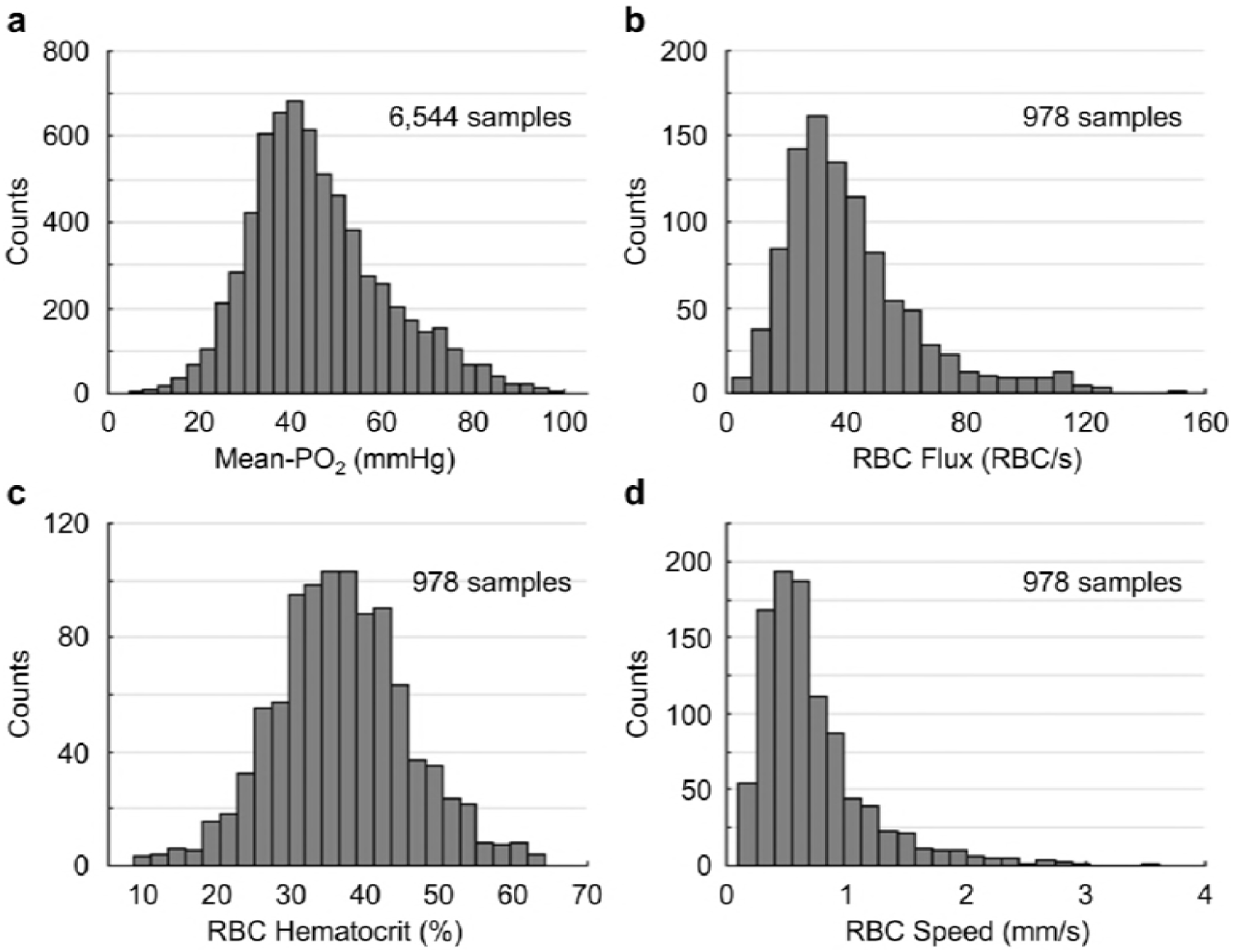
Histograms of capillary Mean-PO_2_, RBC flux, hematocrit and speed. **a.** Capillary Mean-PO_2_ was measured in 6,544 capillary segments, across n = 15 mice. The Mean-PO_2_ range is from ∼5 mmHg to ∼100 mmHg (mean value: 45.6±1.4 mmHg). **b-d.** Capillary RBC flux, hematocrit and speed were measured in a subset of capillaries (967 segments). The RBC flux range (b) is from ∼2 RBC/s to ∼154 RBC/s (mean value: 41.5±1.2 RBC/s); the hematocrit range (c) is from 8.6 % to 64.4 % (mean value: 37.2±0.7 %); the speed range (d) is from 0.11 mm/s to 3.63 mm/s (mean value: 0.71±0.03 mm/s).

**Supplementary Figure 3.**
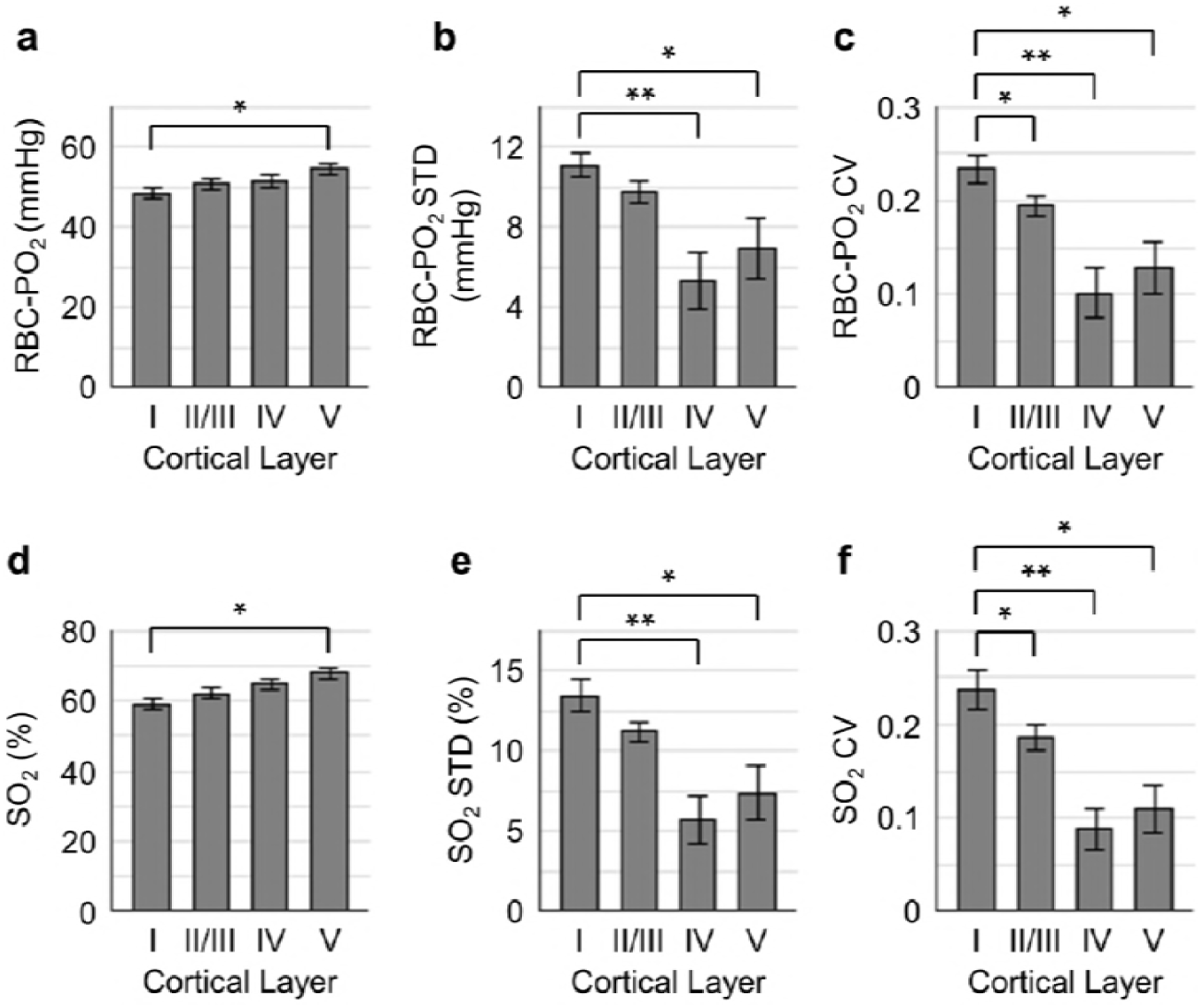
Distributions of capillary RBC-PO_2_, SO_2_ and their STDs and CVs as a function of cortical layer. **a-c.** Distributions of the absolute capillary RBC-PO_2_, RBC-PO_2_ STD and CV as a function of cortical layer, respectively. **d-f.** Distributions of the absolute capillary SO_2_, SO_2_ STD and CV as a function of cortical layer, respectively. The analysis in a-f was made with the measurements from 400, 356, 118 and 104 capillary segments measured in cortical layers I, II/III, IV and V, respectively, across n = 15 mice. Data are expressed as mean±SEM. The single (*) and double (**) asterisk symbols indicate significant differences (Student’s t-test) with P<0.05 and P<0.001, respectively.

**Supplementary Figure 4.**
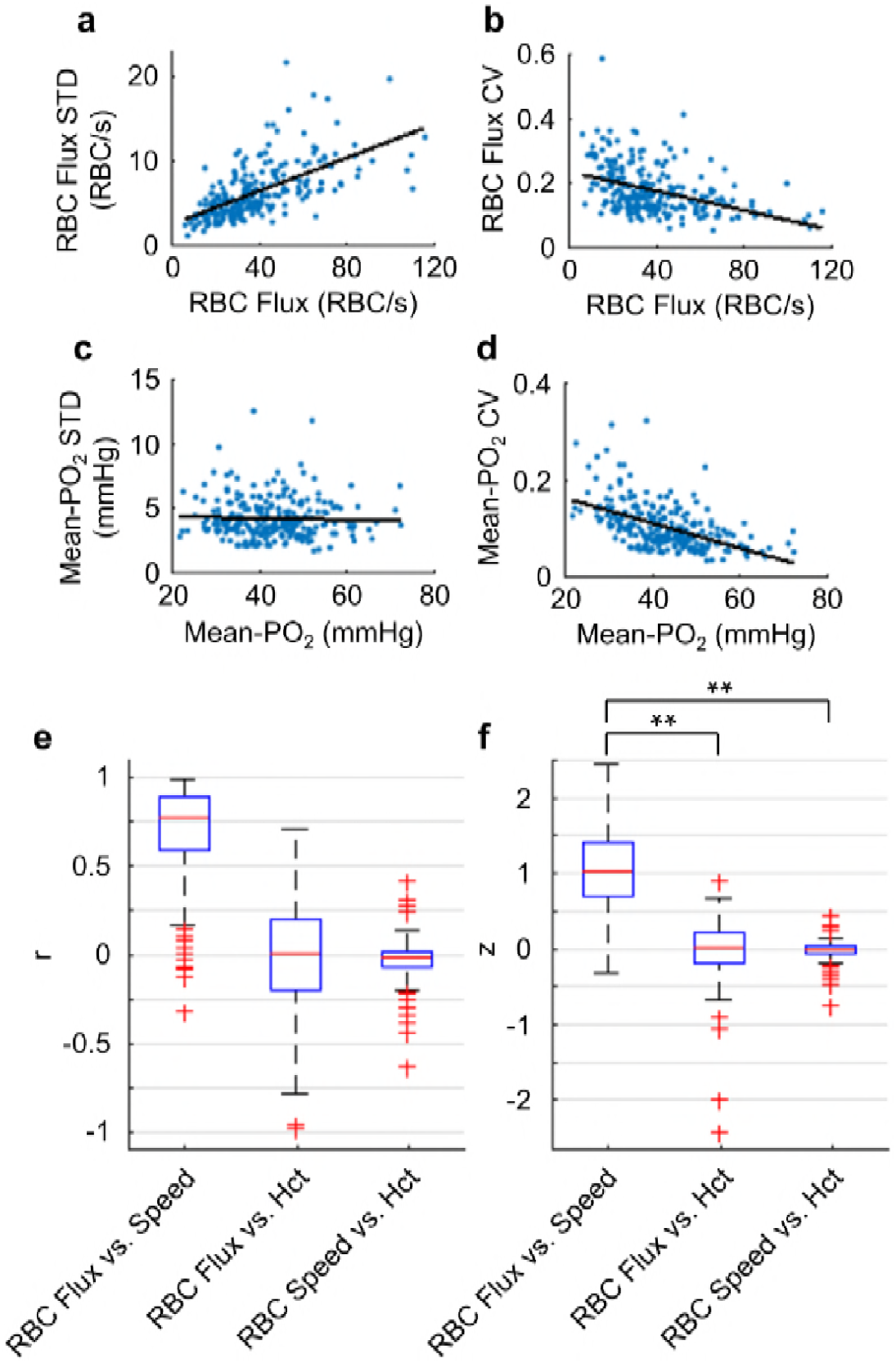
Quantifications of the temporal fluctuations of capillary RBC flux, speed, hematocrit and Mean-PO_2_. **a-d.** Correlations between the STDs and CVs of the temporal fluctuations of capillary RBC flux and Mean-PO_2_ and their mean absolute values, calculated based on the 9-s-long measurements. The data points are represented by the blue dots. The linear regression lines of the correlations are in black. The computed slope (and R^2^) values of the linear regressions in panels a-d are: 0.1 (R^2^ = 0.4), -0.002 s·RBC^-1^ (R^2^ = 0.17), -0.01 (R^2^ = 0.001) and -0.003 mmHg^-1^ (R^2^ = 0.3), respectively. **e and f**. Pairwise correlations between the temporal fluctuations of RBC flux, speed and hematocrit (Hct). The panel e presents the correlation coefficients (r); the panel f presents the Fisher z values. The analysis in **a-f** was made with the measurements from 373 capillary segments in cortical layers I-V, across n = 7 mice. The double asterisk symbol (**) indicates significant difference (Student’s t-test, P<0.001).

**Supplementary Figure 5.**
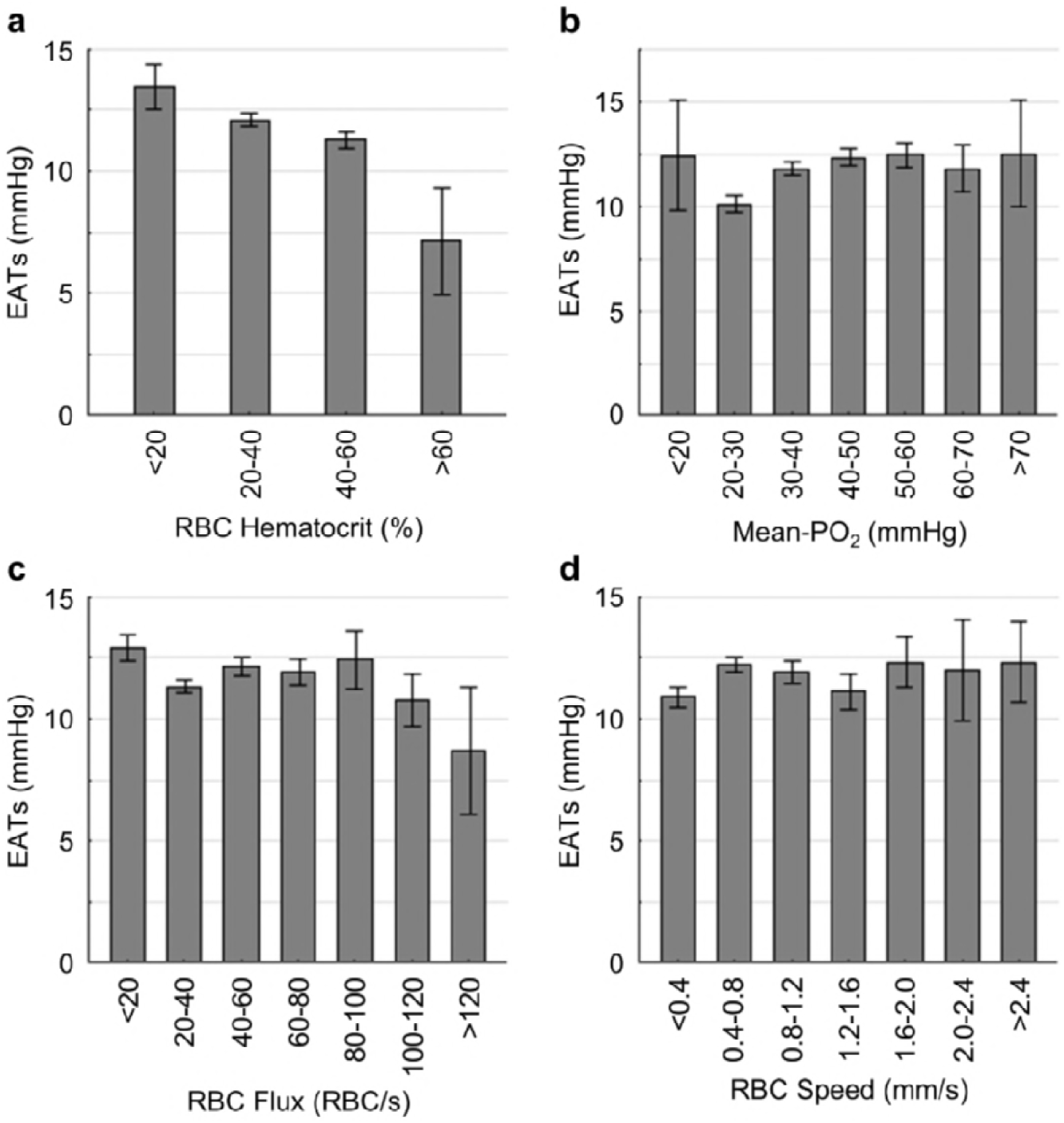
Relations between capillary RBC hematocrit, Mean-PO_2_, flux, speed and EATs. The analysis in a-d was made with the measurements from 373 capillary segments collected in cortical layers I-V, across n = 7 mice. Data are expressed as mean±SEM.

**Supplementary Figure 6.**
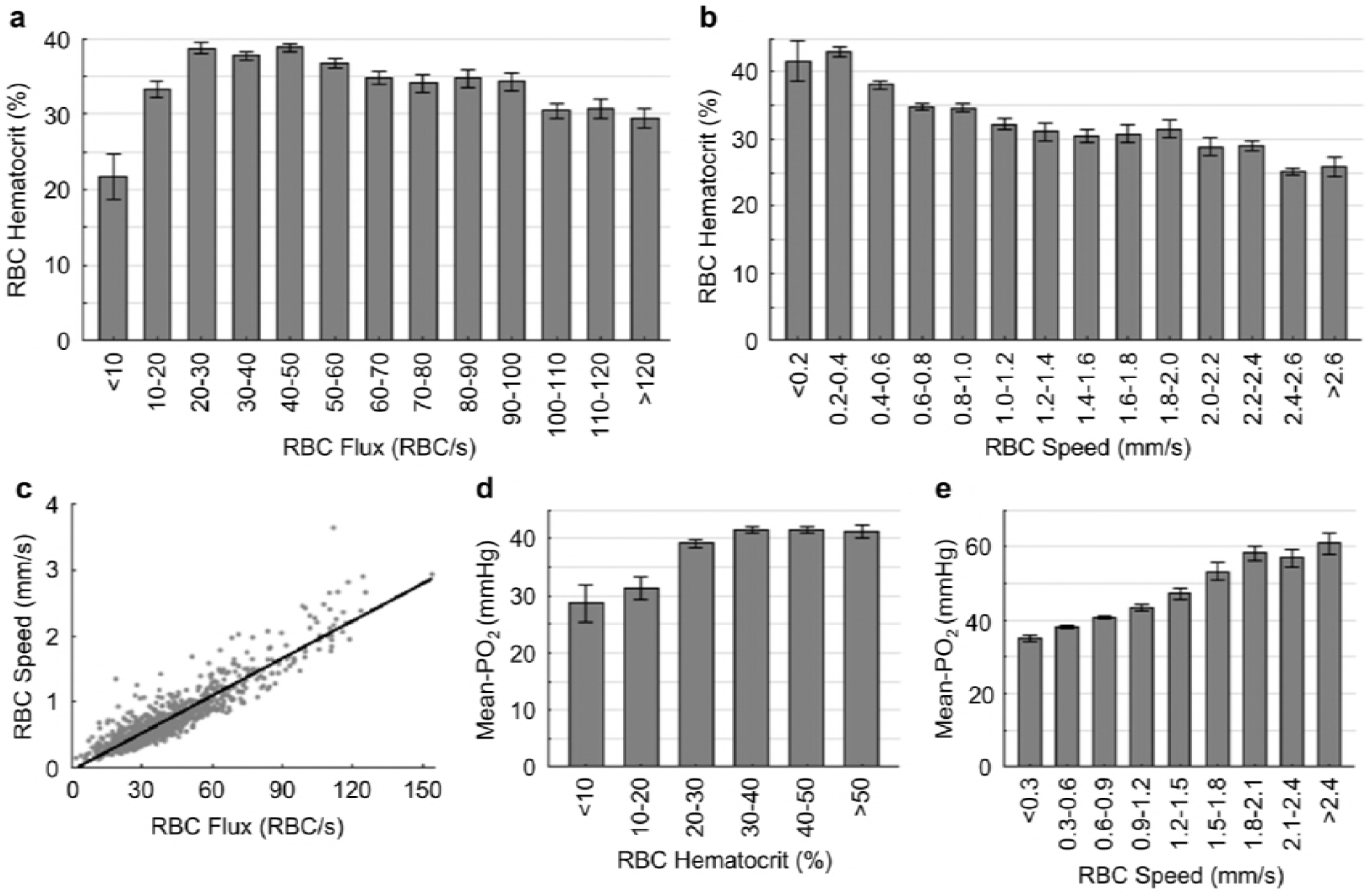
Pairwise relations between capillary RBC flux, speed, hematocrit and Mean-PO_2_. The analysis in a-e was made with the measurements from 978 capillary segments collected in cortical layers I-V, across n = 15 mice. Data are expressed as mean±SEM.

**Supplementary Figure 7.**
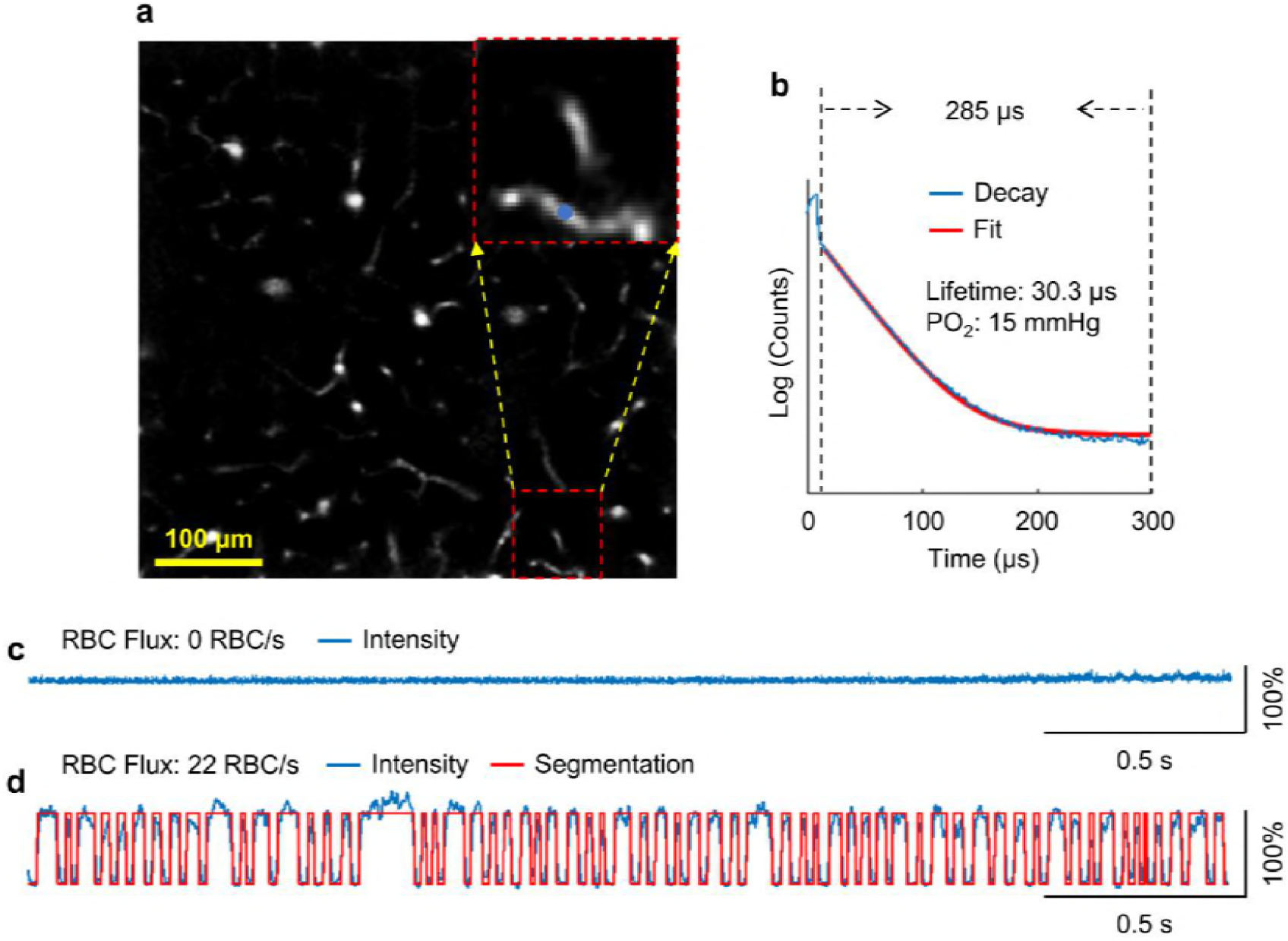
Identification of a capillary segment having stalled RBC flow. **a.** A Sulforhodamine-B labelled mouse cortical microvasculature. The enlarged image includes the stalled capillary segment. The PO_2_ measurement was performed on the location denoted by the blue dot. **b.** The average phosphorescence decay recorded on the measurement location in **a**. The 285-μs-long phosphorescence decay was used to calculate the phosphorescence lifetime. **c.** The associated phosphorescence intensity time course (3-s trace) acquired on the measurement location in **a**. **d.** A phosphorescence intensity time course (3-s trace) acquired in an arbitrary non-stalled capillary segment as a comparison.

**Supplementary Figure 8.**
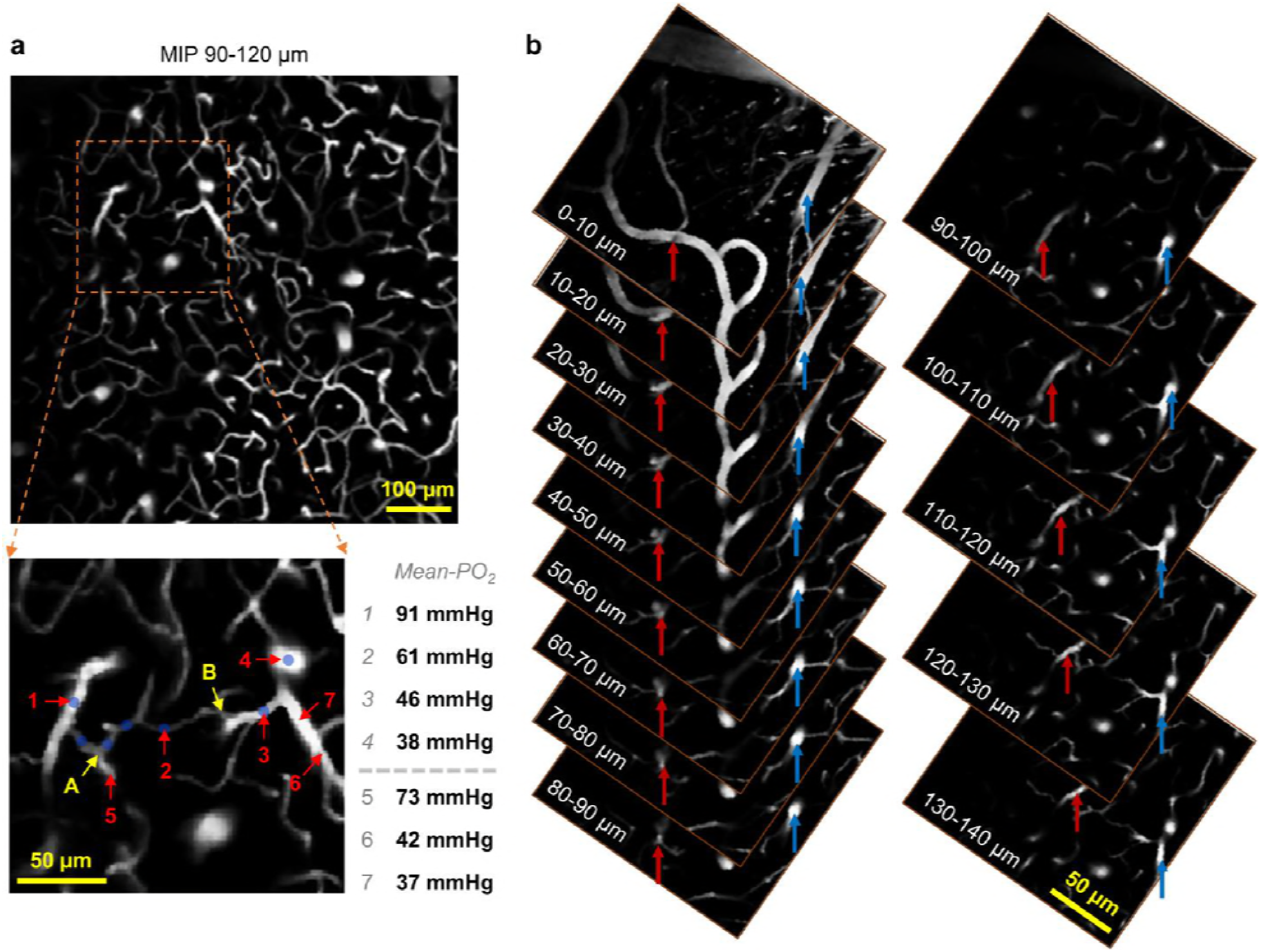
Identification of a suspected thoroughfare capillary. **a.** Upper panel: The maximum intensity projection of the vasculature stack (90-120 μm under cortical surface). Lower panel: The enlarged image includes the vascular paths of the suspected thoroughfare capillary. PO_2_ measurements were performed on the segments labelled with numbers, and their values are shown next to the vascular image. **b.** Tracking of the arteriole (#1 in the lower panel in a) indicated by the red arrow and the venule (#4 in the lower panel in a) indicated by the blue arrow, from the cortical depth of 140 μm to the cortical surface.

The maximum intensity projection of the Sulforhodamine-B labelled microvascular stack (cortical depth: 90-120 μm) is shown in the upper panel of Suppl. Fig. 8a. The enlarged image (Suppl. Fig. 8a, lower panel) includes the vascular paths of the suspected thoroughfare capillary.

The vessel types of the vascular segments #1 and #4 were identified as arteriole and venule, respectively, based on their PO_2_ values and the morphologies of their parent vessels by tracking them with the three-dimensional angiogram to the cortical surface. The complete vascular path, starting from the arteriole (#1) to the venule (#4), consists of the vascular segments marked by the blue dots. PO_2_ measurements were performed on the locations labelled with numbers, and their PO_2_ values are shown at right. The three vascular segments from A to B were identified as capillaries, based on their diameters. The Mean-PO_2_ of the pre-venule capillary (#2) was calculated to be 61 mmHg, much higher than the average capillary Mean-PO_2_ (45.6±1.4 mmHg; Suppl. Fig. 2a). In addition, the vascular path from A to B consists of only three segments; and the physical length from A to B was estimated to be 85 μm. Therefore, this vascular path (A-B) is short in both number of capillary segments and physical length. We thus suspect that the capillary segment #2 is a thoroughfare channel that transported the highly oxygenated blood back to the venule, contributing to the increase in the venous oxygenation towards brain surface.

## REFERENCES

Aaslid, R., Lindegaard, K.F., Sorteberg, W., and Nornes, H. (1989). Cerebral autoregulation dynamics in humans. Stroke 20, 45–52.

Acker, H., and Lübbers, D.W. (1977). The kinetics of local tissue PO2 decrease after perfusion stop within the carotid body of the cat in vivo and in vitro. Pflugers Arch. 369, 135–140.

Alkire, M.T., Pomfrett, C.J.D., Haier, R.J., Gianzero, M.V., Chan, C.M., Jacobsen, B.P., and Fallon, J.H. (1999). Functional Brain Imaging during Anesthesia in Humans Effects of Halothane on Global and Regional Cerebral Glucose Metabolism. Anesthesiol. J. Am. Soc. Anesthesiol. 90, 701–709.

Anenberg, E., Chan, A.W., Xie, Y., LeDue, J.M., and Murphy, T.H. (2015). Optogenetic stimulation of GABA neurons can decrease local neuronal activity while increasing cortical blood flow. J. Cereb. Blood Flow Metab. 35, 1579–1586.

Attwell, D., and Laughlin, S.B. (2001). An Energy Budget for Signaling in the Grey Matter of the Brain. J. Cereb. Blood Flow Metab. 21, 1133–1145.

Attwell, D., Mishra, A., Hall, C.N., O’Farrell, F.M., and Dalkara, T. (2016). What is a pericyte? J. Cereb. Blood Flow Metab. 36, 451–455.

Barker, M.C., Golub, A.S., and Pittman, R.N. (2007). Erythrocyte-associated transients in capillary PO2: an isovolemic hemodilution study in the rat spinotrapezius muscle. Am. J. Physiol. Heart Circ. Physiol. 292, H2540–2549.

Benesch, R.E., Benesch, R., and Yu, C.I. (1969). Oxygenation of hemoglobin in the presence of 2,3-diphosphoglycerate. Effect of temperature, pH, ionic strength, and hemoglobin concentration. Biochemistry 8, 2567–2571.

Berthiaume, A.-A., Hartmann, D.A., Majesky, M.W., Bhat, N.R., and Shih, A.Y. (2018). Pericyte Structural Remodeling in Cerebrovascular Health and Homeostasis. Front. Aging Neurosci. 10.

Blinder, P., Tsai, P.S., Kaufhold, J.P., Knutsen, P.M., Suhl, H., and Kleinfeld, D. (2013). The cortical angiome: an interconnected vascular network with noncolumnar patterns of blood flow. Nat. Neurosci. 16, 889–897.

Blockley, N.P., Griffeth, V.E.M., Stone, A.J., Hare, H.V., and Bulte, D.P. (2015). Sources of systematic error in calibrated BOLD based mapping of baseline oxygen extraction fraction. NeuroImage 122, 105–113.

Braun, R.D., Lanzen, J.L., and Dewhirst, M.W. (1999). Fourier analysis of fluctuations of oxygen tension and blood flow in R3230Ac tumors and muscle in rats. Am. J. Physiol.-Heart Circ. Physiol. 277, H551–H568.

Cai, C., Fordsmann, J.C., Jensen, S.H., Gesslein, B., Lønstrup, M., Hald, B.O., Zambach, S.A., Brodin, B., and Lauritzen, M.J. (2018). Stimulation-induced increases in cerebral blood flow and local capillary vasoconstriction depend on conducted vascular responses. Proc. Natl. Acad. Sci. 115, E5796–E5804.

Cao, R., Li, J., Ning, B., Sun, N., Wang, T., Zuo, Z., and Hu, S. (2017). Functional and oxygen-metabolic photoacoustic microscopy of the awake mouse brain. NeuroImage 150, 77–87.

Chong, S.P., Merkle, C.W., Leahy, C., Radhakrishnan, H., and Srinivasan, V.J. (2015a). Quantitative microvascular hemoglobin mapping using visible light spectroscopic Optical Coherence Tomography. Biomed. Opt. Express 6, 1429–1450.

Chong, S.P., Merkle, C.W., Leahy, C., and Srinivasan, V.J. (2015b). Cerebral metabolic rate of oxygen (CMRO_2_) assessed by combined Doppler and spectroscopic OCT. Biomed. Opt. Express 6, 3941–3951.

Croughwell, N., Smith, L.R., Quill, T., Newman, M., Greeley, W., Kern, F., Lu, J., and Reves, J.G. (1992). The effect of temperature on cerebral metabolism and blood flow in adults during cardiopulmonary bypass. J. Thorac. Cardiovasc. Surg. 103, 549–554.

De Kock C. P. J., Bruno R. M., Spors H., and Sakmann B. (2007). Layer- and cell-type-specific suprathreshold stimulus representation in rat primary somatosensory cortex. J. Physiol. 581, 139–154.

Desjardins, M., Berti, R., Lefebvre, J., Dubeau, S., and Lesage, F. (2014). Aging-related differences in cerebral capillary blood flow in anesthetized rats. Neurobiol. Aging 35, 1947–1955.

Devor, A., Sakadzic, S., Saisan, P.A., Yaseen, M.A., Roussakis, E., Srinivasan, V.J., Vinogradov, S.A., Rosen, B.R., Buxton, R.B., Dale, A.M., et al. (2011). “Overshoot” of O_2_ is required to maintain baseline tissue oxygenation at locations distal to blood vessels. J. Neurosci. Off. J. Soc. Neurosci. 31, 13676–13681.

Diamond, S.G., Huppert, T.J., Kolehmainen, V., Franceschini, M.A., Kaipio, J.P., Arridge, S.R., and Boas, D.A. (2006). Dynamic physiological modeling for functional diffuse optical tomography. NeuroImage 30, 88–101.

Erdener, Ş.E., Tang, J., Sajjadi, A., Kιlιç, K., Kura, S., Schaffer, C.B., and Boas, D.A. (2017). Spatio-temporal dynamics of cerebral capillary segments with stalling red blood cells, Spatio-temporal dynamics of cerebral capillary segments with stalling red blood cells. J. Cereb. Blood Flow Metab. 0271678X17743877.

Esipova, T.V., Rivera-Jacquez, H.J., Weber, B., Masunov, A.E., and Vinogradov, S.A. (2017). Stabilizing g-States in Centrosymmetric Tetrapyrroles: Two-Photon-Absorbing Porphyrins with Bright Phosphorescence. J. Phys. Chem. A 121, 6243–6255.

Eskildsen, S.F., Gyldensted, L., Nagenthiraja, K., Nielsen, R.B., Hansen, M.B., Dalby, R.B., Frandsen, J., Rodell, A., Gyldensted, C., Jespersen, S.N., et al. (2017). Increased cortical capillary transit time heterogeneity in Alzheimer’s disease: a DSC-MRI perfusion study. Neurobiol. Aging 50, 107–118.

Finikova, O.S., Lebedev, A.Y., Aprelev, A., Troxler, T., Gao, F., Garnacho, C., Muro, S., Hochstrasser, R.M., and Vinogradov, S.A. (2008). Oxygen Microscopy by Two-Photon-Excited Phosphorescence. ChemPhysChem 9, 1673–1679.

Gagnon, L., Smith, A.F., Boas, D.A., Devor, A., Secomb, T.W., and Sakadžić, S. (2016). Modeling of Cerebral Oxygen Transport Based on In vivo Microscopic Imaging of Microvascular Network Structure, Blood Flow, and Oxygenation. Front. Comput. Neurosci. 10, 82.

Girouard, H., and Iadecola, C. (2006). Neurovascular coupling in the normal brain and in hypertension, stroke, and Alzheimer disease. J. Appl. Physiol. Bethesda Md 1985 100, 328–335.

Goldberg, N.D., Passonneau, J.V., and Lowry, O.H. (1966). Effects of Changes in Brain Metabolism on the Levels of Citric Acid Cycle Intermediates. J. Biol. Chem. 241, 3997–4003.

Goldey, G.J., Roumis, D.K., Glickfeld, L.L., Kerlin, A.M., Reid, R.C., Bonin, V., Schafer, D.P., and Andermann, M.L. (2014). Removable cranial windows for long-term imaging in awake mice. Nat. Protoc. 9, 2515–2538.

Golub, A.S., and Pittman, R.N. (2005). Erythrocyte-associated transients in PO2 revealed in capillaries of rat mesentery. Am. J. Physiol. Heart Circ. Physiol. 288, H2735–2743.

Griffeth, V.E.M., and Buxton, R.B. (2011). A theoretical framework for estimating cerebral oxygen metabolism changes using the calibrated-BOLD method: Modeling the effects of blood volume distribution, hematocrit, oxygen extraction fraction, and tissue signal properties on the BOLD signal. NeuroImage 58, 198–212.

Gutiérrez-Jiménez, E., Cai, C., Mikkelsen, I.K., Rasmussen, P.M., Angleys, H., Merrild, M., Mouridsen, K., Jespersen, S.N., Lee, J., Iversen, N.K., et al. (2016). Effect of electrical forepaw stimulation on capillary transit-time heterogeneity (CTH). J. Cereb. Blood Flow Metab. 36, 2072–2086.

Gutiérrez-Jiménez, E., Angleys, H., Rasmussen, P.M., West, M.J., Catalini, L., Iversen, N.K., Jensen, M.S., Frische, S., and Østergaard, L. (2018). Disturbances in the control of capillary flow in an aged APPswe/PS1ΔE9 model of Alzheimer’s disease. Neurobiol. Aging 62, 82–94.

Hall, C.N., Reynell, C., Gesslein, B., Hamilton, N.B., Mishra, A., Sutherland, B.A., O’Farrell, F.M., Buchan, A.M., Lauritzen, M., and Attwell, D. (2014). Capillary pericytes regulate cerebral blood flow in health and disease. Nature 508, 55–60.

Hanks, J.H., and Wallace, R.E. (1949). Relation of Oxygen and Temperature in the Preservation of Tissues by Refrigeration, Relation of Oxygen and Temperature in the Preservation of Tissues by Refrigeration. Proc. Soc. Exp. Biol. Med. 71, 196–200.

Hartmann, D.A., Underly, R.G., Grant, R.I., Watson, A.N., Lindner, V., and Shih, A.Y. (2015). Pericyte structure and distribution in the cerebral cortex revealed by high-resolution imaging of transgenic mice. Neurophotonics 2, 041402.

Hellums, J.D. (1977). The resistance to oxygen transport in the capillaries relative to that in the surrounding tissue. Microvasc. Res. 13, 131–136.

Hernandez, J.C.C., Kersbergen, C., Muse, V., Ivasyk, I., Bracko, O., Haft-Javaherian, M., Kang, Y., Zhou, J., Beverly, J.D., Slack, E., et al. (2016). STALLED BLOOD FLOW IN BRAIN CAPILLARIES IS RESPONSIBLE FOR REDUCED CORTICAL PERFUSION IN A MOUSE MODEL OF ALZHEIMER’S DISEASE. Alzheimers Dement. J. Alzheimers Assoc. 12, P1049–P1050.

Hill, R.A., Tong, L., Yuan, P., Murikinati, S., Gupta, S., and Grutzendler, J. (2015). Regional Blood Flow in the Normal and Ischemic Brain Is Controlled by Arteriolar Smooth Muscle Cell Contractility and Not by Capillary Pericytes. Neuron 87, 95–110.

Hu, S., Maslov, K.I., Tsytsarev, V., and Wang, L.V. (2009). Functional transcranial brain imaging by optical-resolution photoacoustic microscopy. J. Biomed. Opt. 14, 040503.

Hudetz, A.G., Fehér, G., and Kampine, J.P. (1996). Heterogeneous Autoregulation of Cerebrocortical Capillary Flow: Evidence for Functional Thoroughfare Channels? Microvasc. Res. 51, 131–136.

Hyder, F., Rothman, D.L., and Bennett, M.R. (2013). Cortical energy demands of signaling and nonsignaling components in brain are conserved across mammalian species and activity levels. Proc. Natl. Acad. Sci. 110, 3549–3554.

Iadecola, C. (2016). Vascular and Metabolic Factors in Alzheimer’s Disease and Related Dementias: Introduction. Cell. Mol. Neurobiol. 36, 151–154.

Iadecola, C. (2017). The Neurovascular Unit Coming of Age: A Journey through Neurovascular Coupling in Health and Disease. Neuron 96, 17–42.

Iordanova, B., Vazquez, A.L., Poplawsky, A.J., Fukuda, M., and Kim, S.-G. (2015). Neural and Hemodynamic Responses to Optogenetic and Sensory Stimulation in the Rat Somatosensory Cortex. J. Cereb. Blood Flow Metab. 35, 922–932.

Jespersen, S.N., and Østergaard, L. (2012). The roles of cerebral blood flow, capillary transit time heterogeneity, and oxygen tension in brain oxygenation and metabolism. J. Cereb. Blood Flow Metab. 32, 264–277.

Jones, P.F., and Siegel, S. (1969). Temperature Effects on the Phosphorescence of Aromatic Hydrocarbons in Poly(methylmethacrylate). J. Chem. Phys. 50, 1134–1140.

Kazmi, S.M.S., Salvaggio, A.J., Estrada, A.D., Hemati, M.A., Shaydyuk, N.K., Roussakis, E., Jones, T.A., Vinogradov, S.A., and Dunn, A.K. (2013). Three-dimensional mapping of oxygen tension in cortical arterioles before and after occlusion. Biomed. Opt. Express 4, 1061–1073.

Kimura, H., Braun, R.D., Ong, E.T., Hsu, R., Secomb, T.W., Papahadjopoulos, D., Hong, K., and Dewhirst, M.W. (1996). Fluctuations in Red Cell Flux in Tumor Microvessels Can Lead to Transient Hypoxia and Reoxygenation in Tumor Parenchyma. Cancer Res. 56, 5522–5528.

Kisler, K., Nelson, A.R., Rege, S.V., Ramanathan, A., Wang, Y., Ahuja, A., Lazic, D., Tsai, P.S., Zhao, Z., Zhou, Y., et al. (2017). Pericyte degeneration leads to neurovascular uncoupling and limits oxygen supply to brain. Nat. Neurosci. 20, 406–416.

Kleinfeld, D., Mitra, P.P., Helmchen, F., and Denk, W. (1998). Fluctuations and stimulus-induced changes in blood flow observed in individual capillaries in layers 2 through 4 of rat neocortex. Proc. Natl. Acad. Sci. 95, 15741–15746.

Komiyama, T., Sato, T.R., O’Connor, D.H., Zhang, Y.-X., Huber, D., Hooks, B.M., Gabitto, M., and Svoboda, K. (2010). Learning-related fine-scale specificity imaged in motor cortex circuits of behaving mice. Nature 464, 1182–1186.

Land, P.W., and Simons, D.J. Cytochrome oxidase staining in the rat smI barrel cortex. J. Comp. Neurol. 238, 225–235.

Lebedev, A.Y., Cheprakov, A.V., Sakadžić, S., Boas, D.A., Wilson, D.F., and Vinogradov, S.A. (2009). Dendritic Phosphorescent Probes for Oxygen Imaging in Biological Systems. ACS Appl. Mater. Interfaces 1, 1292–1304.

Lecoq, J., Tiret, P., Najac, M., Shepherd, G.M., Greer, C.A., and Charpak, S. (2009). Odor-Evoked Oxygen Consumption by Action Potential and Synaptic Transmission in the Olfactory Bulb. J. Neurosci. 29, 1424–1433.

Lecoq, J., Parpaleix, A., Roussakis, E., Ducros, M., Houssen, Y.G., Vinogradov, S.A., and Charpak, S. (2011). Simultaneous two-photon imaging of oxygen and blood flow in deep cerebral vessels. Nat. Med. 17, 893–898.

Lee, J., Wu, W., and Boas, D.A. (2015). Early capillary flux homogenization in response to neural activation. J. Cereb. Blood Flow Metab. 0271678X15605851.

Lefort, S., Tomm, C., Floyd Sarria, J.-C., and Petersen, C.C.H. (2009). The excitatory neuronal network of the C2 barrel column in mouse primary somatosensory cortex. Neuron 61, 301–316.

Lücker, A., Secomb, T.W., Weber, B., and Jenny, P. (2017). The relative influence of hematocrit and red blood cell velocity on oxygen transport from capillaries to tissue. Microcirculation 24, n/a–n/a.

Lücker, A., Secomb, T.W., Weber, B., and Jenny, P. (2018). The Relation Between Capillary Transit Times and Hemoglobin Saturation Heterogeneity. Part 1: Theoretical Models. Front. Physiol. 9.

Lyons, D.G., Parpaleix, A., Roche, M., and Charpak, S. (2016). Mapping oxygen concentration in the awake mouse brain. ELife 5, e12024.

Mateo, C., Avermann, M., Gentet, L.J., Zhang, F., Deisseroth, K., and Petersen, C.C.H. (2011). In Vivo Optogenetic Stimulation of Neocortical Excitatory Neurons Drives Brain-State-Dependent Inhibition. Curr. Biol. 21, 1593–1602.

Merkle, C.W., and Srinivasan, V.J. (2016). Laminar microvascular transit time distribution in the mouse somatosensory cortex revealed by Dynamic Contrast Optical Coherence Tomography. NeuroImage 125, 350–362.

Mintun, M.A., Lundstrom, B.N., Snyder, A.Z., Vlassenko, A.G., Shulman, G.L., and Raichle, M.E. (2001). Blood flow and oxygen delivery to human brain during functional activity: Theoretical modeling and experimental data. Proc. Natl. Acad. Sci. 98, 6859–6864.

Moeini, M., Lu, X., Avti, P.K., Damseh, R., Bélanger, S., Picard, F., Boas, D., Kakkar, A., and Lesage, F. (2018). Compromised microvascular oxygen delivery increases brain tissue vulnerability with age. Sci. Rep. 8, 8219.

Momjian-Mayor, I., and Baron, J.-C. (2005). The Pathophysiology of Watershed Infarction in Internal Carotid Artery Disease: Review of Cerebral Perfusion Studies. Stroke 36, 567–577.

Müller, K., Courtois, G., Ursini, M.V., and Schwaninger, M. (2017). New Insight Into the Pathogenesis of Cerebral Small-Vessel Diseases. Stroke 48, 520–527.

Nguyen, J., Nishimura, N., Fetcho, R.N., Iadecola, C., and Schaffer, C.B. (2011). Occlusion of Cortical Ascending Venules Causes Blood Flow Decreases, Reversals in Flow Direction, and Vessel Dilation in Upstream Capillaries. J. Cereb. Blood Flow Metab. 31, 2243–2254.

Nielsen, R.B., Egefjord, L., Angleys, H., Mouridsen, K., Gejl, M., Møller, A., Brock, B., Brændgaard, H., Gottrup, H., Rungby, J., et al. (2017). Capillary dysfunction is associated with symptom severity and neurodegeneration in Alzheimer’s disease. Alzheimers Dement. J. Alzheimers Assoc. 13, 1143–1153.

Ob, P., S, S., and L, E. (1990). Cerebral autoregulation. Cerebrovasc. Brain Metab. Rev. 2, 161–192.

Ogawa, S., Lee, T.M., Kay, A.R., and Tank, D.W. (1990). Brain magnetic resonance imaging with contrast dependent on blood oxygenation. Proc. Natl. Acad. Sci. 87, 9868–9872.

Østergaard, L., Aamand, R., Gutiérrez-Jiménez, E., Ho, Y.-C.L., Blicher, J.U., Madsen, S.M., Nagenthiraja, K., Dalby, R.B., Drasbek, K.R., Møller, A., et al. (2013). The capillary dysfunction hypothesis of Alzheimer’s disease. Neurobiol. Aging 34, 1018–1031.

Østergaard, L., Engedal, T.S., Aamand, R., Mikkelsen, R., Iversen, N.K., Anzabi, M., Næss-Schmidt, E.T., Drasbek, K.R., Bay, V., Blicher, J.U., et al. (2014a). Capillary transit time heterogeneity and flow-metabolism coupling after traumatic brain injury. J. Cereb. Blood Flow Metab. 34, 1585–1598.

Østergaard, L., Kristiansen, S.B., Angleys, H., Frøkiær, J., Hasenkam, J.M., Jespersen, S.N., and Bøtker, H.E. (2014b). The role of capillary transit time heterogeneity in myocardial oxygenation and ischemic heart disease. Basic Res. Cardiol. 109, 1–18.

Pantoni, L. (2010). Cerebral small vessel disease: from pathogenesis and clinical characteristics to therapeutic challenges. Lancet Neurol. 9, 689–701.

Parpaleix, A., Houssen, Y.G., and Charpak, S. (2013). Imaging local neuronal activity by monitoring PO2 transients in capillaries. Nat. Med. 19, 241–246.

Patel, U. (1983). Non-random distribution of blood vessels in the posterior region of the rat somatosensory cortex. Brain Res. 289, 65–70.

Peppiatt, C.M., Howarth, C., Mobbs, P., and Attwell, D. (2006). Bidirectional control of CNS capillary diameter by pericytes. Nature 443, 700–704.

Pittman, R.N. (2011). Regulation of Tissue Oxygenation (San Rafael (CA): Morgan & Claypool Life Sciences).

Pray, H.A. (1952). Solubility of Hydrogen, Oxygen, Nitrogen, and Helium in Water. Ind Eng Chem 44, 1146–1151.

Pries, A.R., Secomb, T.W., Gessner, T., Sperandio, M.B., Gross, J.F., and Gaehtgens, P. (1994). Resistance to blood flow in microvessels in vivo. Circ. Res. 75, 904–915.

Pries, A.R., Secomb, T.W., and Gaehtgens, P. (1998). Structural adaptation and stability of microvascular networks: theory and simulations. Am. J. Physiol. 275, H349–360.

Raichle, M.E., and Gusnard, D.A. (2002). Appraising the brain’s energy budget. Proc. Natl. Acad. Sci. 99, 10237–10239.

Raichle, M.E., MacLeod, A.M., Snyder, A.Z., Powers, W.J., Gusnard, D.A., and Shulman, G.L. (2001). A default mode of brain function. Proc. Natl. Acad. Sci. 98, 676–682.

Rasmussen, P.M., Jespersen, S.N., and Østergaard, L. (2015). The effects of transit time heterogeneity on brain oxygenation during rest and functional activation. J. Cereb. Blood Flow Metab. 35, 432–442.

Reglin, B., Secomb, T.W., and Pries, A.R. (2009). Structural adaptation of microvessel diameters in response to metabolic stimuli: where are the oxygen sensors? Am. J. Physiol. - Heart Circ. Physiol. 297, H2206–H2219.

Reglin, B., Secomb, T.W., and Pries, A.R. (2017). Structural Control of Microvessel Diameters: Origins of Metabolic Signals. Front. Physiol. 8, 813.

Rosomoff, H., and Holaday, D. (1954). Cerebral blood flow and cerebral oxygen consumption during hypothermia. Am. J. Physiol. 179, 85–88.

Rossing, R.G., and Cain, S.M. (1966). A nomogram relating pO_2_, pH, temperature, and hemoglobin saturation in the dog. J. Appl. Physiol. 21, 195–201.

Sakadžić, S., Roussakis, E., Yaseen, M.A., Mandeville, E.T., Srinivasan, V.J., Arai, K., Ruvinskaya, S., Devor, A., Lo, E.H., Vinogradov, S.A., et al. (2010). Two-photon high-resolution measurement of partial pressure of oxygen in cerebral vasculature and tissue. Nat. Methods 7, 755–759.

Sakadžić, S., Roussakis, E., Yaseen, M.A., Mandeville, E.T., Srinivasan, V.J., Arai, K., Ruvinskaya, S., Wu, W., Devor, A., Lo, E.H., et al. (2011). Cerebral blood oxygenation measurement based on oxygen-dependent quenching of phosphorescence. J. Vis. Exp. JoVE.

Sakadžić, S., Mandeville, E.T., Gagnon, L., Musacchia, J.J., Yaseen, M.A., Yucel, M.A., Lefebvre, J., Lesage, F., Dale, A.M., Eikermann-Haerter, K., et al. (2014). Large arteriolar component of oxygen delivery implies a safe margin of oxygen supply to cerebral tissue. Nat. Commun. 5.

Sakadžić, S., Lee, J., Boas, D.A., and Ayata, C. (2015). High-resolution in vivo optical imaging of stroke injury and repair. Brain Res. 1623, 174–192.

Sakadžić, S., Yaseen, M.A., Jaswal, R., Roussakis, E., Dale, A.M., Buxton, R.B., Vinogradov, S.A., Boas, D.A., and Devor, A. (2016). Two-photon microscopy measurement of cerebral metabolic rate of oxygen using periarteriolar oxygen concentration gradients. Neurophotonics 3.

Santisakultarm, T.P., Cornelius, N.R., Nishimura, N., Schafer, A.I., Silver, R.T., Doerschuk, P.C., Olbricht, W.L., and Schaffer, C.B. (2012). In vivo two-photon excited fluorescence microscopy reveals cardiac- and respiration-dependent pulsatile blood flow in cortical blood vessels in mice. Am. J. Physiol. Heart Circ. Physiol. 302, H1367–1377.

Santisakultarm, T.P., Paduano, C.Q., Stokol, T., Southard, T.L., Nishimura, N., Skoda, R.C., Olbricht, W.L., Schafer, A.I., Silver, R.T., and Schaffer, C.B. (2014). Stalled cerebral capillary blood flow in mouse models of essential thrombocythemia and polycythemia vera revealed by in vivo two-photon imaging. J. Thromb. Haemost. JTH 12, 2120–2130.

Sato, Y., Nakajima, S., Shiraga, N., Atsumi, H., Yoshida, S., Koller, T., Gerig, G., and Kikinis, R. (1998). Three-dimensional multi-scale line filter for segmentation and visualization of curvilinear structures in medical images. Med. Image Anal. 2, 143–168.

Secomb, T.W., Hsu, R., Beamer, N.B., and Coull, B.M. (2000). Theoretical simulation of oxygen transport to brain by networks of microvessels: effects of oxygen supply and demand on tissue hypoxia. Microcirc. N. Y. N 1994 7, 237–247.

Shirey, M.J., Smith, J.B., Kudlik, D.E., Huo, B.-X., Greene, S.E., and Drew, P.J. (2015). Brief anesthesia, but not voluntary locomotion, significantly alters cortical temperature. J. Neurophysiol. 114, 309–322.

Siero, J.C.W., Petridou, N., Hoogduin, H., Luijten, P.R., and Ramsey, N.F. (2011). Cortical depth-dependent temporal dynamics of the BOLD response in the human brain. J. Cereb. Blood Flow Metab. Off. J. Int. Soc. Cereb. Blood Flow Metab. 31, 1999–2008.

Silva, A.C., and Koretsky, A.P. (2002). Laminar specificity of functional MRI onset times during somatosensory stimulation in rat. Proc. Natl. Acad. Sci. U. S. A. 99, 15182–15187.

Soukup, J., Zauner, A., Doppenberg, E.M.R., Menzel, M., Gilman, C., Young, H.F., and Bullock, R. (2002). The Importance of Brain Temperature in Patients after Severe Head Injury: Relationship to Intracranial Pressure, Cerebral Perfusion Pressure, Cerebral Blood Flow, and Outcome. J. Neurotrauma 19, 559–571.

Srinivasan, V.J., Yu, E., Radhakrishnan, H., Can, A., Climov, M., Leahy, C., Ayata, C., and Eikermann-Haerter, K. (2015). Micro-heterogeneity of flow in a mouse model of chronic cerebral hypoperfusion revealed by longitudinal Doppler optical coherence tomography and angiography. J. Cereb. Blood Flow Metab. Off. J. Int. Soc. Cereb. Blood Flow Metab. 35, 1552–1560.

Stefanovic, B., Hutchinson, E., Yakovleva, V., Schram, V., Russell, J.T., Belluscio, L., Koretsky, A.P., and Silva, A.C. (2007). Functional reactivity of cerebral capillaries. J. Cereb. Blood Flow Metab. 28, 961–972.

Uchida, K., Reilly, M.P., and Asakura, T. (1998). Molecular Stability and Function of Mouse Hemoglobins. Zoolog. Sci. 15, 703–706.

Unekawa, M., Tomita, M., Tomita, Y., Toriumi, H., Miyaki, K., and Suzuki, N. (2010). RBC velocities in single capillaries of mouse and rat brains are the same, despite 10-fold difference in body size. Brain Res. 1320, 69–73.

Vanderkooi, J.M., Maniara, G., Green, T.J., and Wilson, D.F. (1987). An optical method for measurement of dioxygen concentration based upon quenching of phosphorescence. J. Biol. Chem. 262, 5476–5482.

VanTeeffelen, J.W.G.E., Constantinescu, A.A., Brands, J., Spaan, J.A.E., and Vink, H. Bradykinin- and sodium nitroprusside-induced increases in capillary tube haematocrit in mouse cremaster muscle are associated with impaired glycocalyx barrier properties. J. Physiol. 586, 3207–3218.

Vazquez, A.L., Cohen, E.R., Gulani, V., Hernandez-Garcia, L., Zheng, Y., Lee, G.R., Kim, S.-G., Grotberg, J.B., and Noll, D.C. (2006). Vascular dynamics and BOLD fMRI: CBF level effects and analysis considerations. NeuroImage 32, 1642–1655.

Vazquez, A.L., Masamoto, K., Fukuda, M., and Kim, S.-G. (2010). Cerebral oxygen delivery and consumption during evoked neural activity. Front. Neuroenergetics 2.

Wang, Y., Hu, S., Maslov, K., Zhang, Y., Xia, Y., and Wang, L.V. (2011). In vivo integrated photoacoustic and confocal microscopy of hemoglobin oxygen saturation and oxygen partial pressure. Opt. Lett. 36, 1029–1031.

Wardlaw, J.M., Smith, C., and Dichgans, M. (2013). Mechanisms of sporadic cerebral small vessel disease: insights from neuroimaging. Lancet Neurol. 12, 483–497.

Whalen, W.J., and Nair, P. (1975). Some factors affecting tissue Po2 in the carotid body. J. Appl. Physiol. 39, 562–566.

Wu, J., Guo, C., Chen, S., Jiang, T., He, Y., Ding, W., Yang, Z., Luo, Q., and Gong, H. (2016). Direct 3D Analyses Reveal Barrel-Specific Vascular Distribution and Cross-Barrel Branching in the Mouse Barrel Cortex. Cereb. Cortex 26, 23–31.

Yaseen, M.A., Srinivasan, V.J., Sakadžić, S., Wu, W., Ruvinskaya, S., Vinogradov, S.A., and Boas, D. (2009). Optical monitoring of oxygen tension in cortical microvessels with confocal microscopy. Opt. Express 17, 22341–22350.

Yaseen, M.A., Srinivasan, V.J., Gorczynska, I., Fujimoto, J.G., Boas, D.A., and Sakadžić, S. (2015). Multimodal optical imaging system for in vivo investigation of cerebral oxygen delivery and energy metabolism. Biomed. Opt. Express 6, 4994–5007.

Yu, X., Qian, C., Chen, D., Dodd, S.J., and Koretsky, A.P. (2014). Deciphering laminar-specific neural inputs with line-scanning fMRI. Nat. Methods 11, 55–58.

Zlokovic, B.V. (2011). Neurovascular pathways to neurodegeneration in Alzheimer’s disease and other disorders. Nat. Rev. Neurosci. 12, 723–738.

